# Mechanism of R-loop formation and conformational activation of Cas9

**DOI:** 10.1101/2021.09.16.460614

**Authors:** Martin Pacesa, Luuk Loeff, Irma Querques, Lena M. Muckenfuss, Marta Sawicka, Martin Jinek

## Abstract

Cas9 is a CRISPR-associated endonuclease capable of RNA-guided, site-specific DNA cleavage^1–3^. The programmable activity of Cas9 has been widely utilized for genome editing applications^4–6^, yet its precise mechanisms of target DNA binding and off-target discrimination remain incompletely understood. Here we report a series of cryo-EM structures of *Streptococcus pyogenes* Cas9 capturing the directional process of target DNA hybridization. In the early phase of R-loop formation, the Cas9 REC2 and REC3 domains form a positively charged cleft that accommodates the distal end of the target DNA duplex. Guide-target hybridization past the seed region induces rearrangements of the REC2/3 domains and relocation of the HNH nuclease domain to assume a catalytically incompetent checkpoint conformation. Completion of the guide-target heteroduplex triggers conformational activation of the HNH nuclease domain, enabled by distortion of the guide-target heteroduplex, and complementary REC2/3 domain rearrangements. Together, these results establish a structural framework for target DNA-dependent activation of Cas9 that sheds light on its conformational checkpoint mechanism and may facilitate the development of novel Cas9 variants and guide RNA designs with enhanced specificity and activity.

## Main

Cas9 enzymes rely on a dual guide RNA structure consisting of a CRISPR RNA (crRNA) guide and a trans-activating CRISPR RNA (tracrRNA) coactivator to cleave complementary DNA targets. *Streptococcus pyogenes* Cas9 (SpCas9) has found widespread use as a programmable DNA targeting tool in genome editing and gene targeting applications^4–6^. Target DNA binding by SpCas9 is dependent on initial recognition of an NGG protospacer-adjacent motif (PAM) downstream of the target site^2,7–9^, which triggers local DNA strand separation to initiate its directional hybridization with a 20-nt segment in the guide crRNA to form an R-loop structure^7,10,11^. Target strand binding is facilitated by structural pre-ordering of nucleotides 11-20 of the crRNA (counting from the 5’ end), termed the seed sequence, in an A form-like conformation^8,12^. Formation of a complete R-loop leads to the activation of the Cas9 HNH and RuvC nuclease domains to catalyse cleavage of the target (TS) and non-target (NTS) DNA strands, respectively^2,8,13^. Although highly specific, SpCas9 cleaves off-target sites with imperfect complementarity to the guide RNA, often resulting in considerable levels of off-target genome editing^14–18^. The off-target activity is dependent on the number, type, and positioning of base mismatches within the guide-target heteroduplex^15,19–21^. PAM-proximal mismatches within the seed region are discriminated against by substantially increased dissociation rates^11,19,21,22^ whereas PAM-distal mismatches are compatible with stable DNA binding^13,19,21,23,24^. Such off-targets are instead discriminated by a conformational checkpoint mechanism that monitors the integrity of the guide-target duplex to induce conformational activation of the nuclease domains^11,19,21,22,13,23,24^. Structural, biophysical and computational studies of SpCas9 have shed light on the mechanism of guide RNA binding, PAM recognition, and nuclease activation, revealing that the enzyme undergoes extensive conformational rearrangements throughout these steps. In particular, high-resolution structures of the fully-bound target DNA complex of SpCas9^25–28^ have revealed a target DNA-dependent conformational rearrangement of the Cas9 REC-lobe that is necessary for cleavage activation. However, the mechanisms that underpin R-loop formation and off-target discrimination during conformational activation have remained elusive.

### Cryo-EM analysis of R-loop formation

To investigate the mechanism of R-loop formation, we initially determined the minimal extent of target DNA complementarity necessary for stable binding using fluorescence-coupled size exclusion chromatography, revealing that the presence of six complementary nucleotides in the PAM-proximal region of the target DNA heteroduplex is sufficient for stable association with the SpCas9-guide RNA complex. (**Extended Data Fig. 1**). Subsequently, catalytically-inactive SpCas9 (dCas9) was reconstituted with a single-molecule guide RNA (sgRNA) and partially-matched DNA targets containing 6, 8, 10, 12, 14, 16 complementary nucleotides upstream of the PAM (**Fig. 1a, Extended Data Fig. 2**). The resulting complexes were analysed by cryo-EM, yielding molecular reconstructions at resolutions of 3.0–4.1 Å (**Extended Data Fig. 3, Extended Data Table 1**). We additionally determined cryo-EM reconstructions of wild-type SpCas9 bound to 18-nt complementary DNA targets in the presence of l mM and 10 mM Mg^2+^, representing the checkpoint and catalytically active states, respectively (**Extended Data Fig. 3, Extended Data Table 1**). 3D variability analysis^29^ was used to analyse conformational heterogeneity within each complex (**Supplementary Videos 1-8**). Most of the detected variability within each complex can be attributed to the PAM-distal duplex and the REC2, REC3, and HNH domains, suggestive of conformational equilibrium sampling. The resulting structural models are representative of the most abundant conformational state of each complex (**Extended Data Fig. 4**).

**Figure 1.**
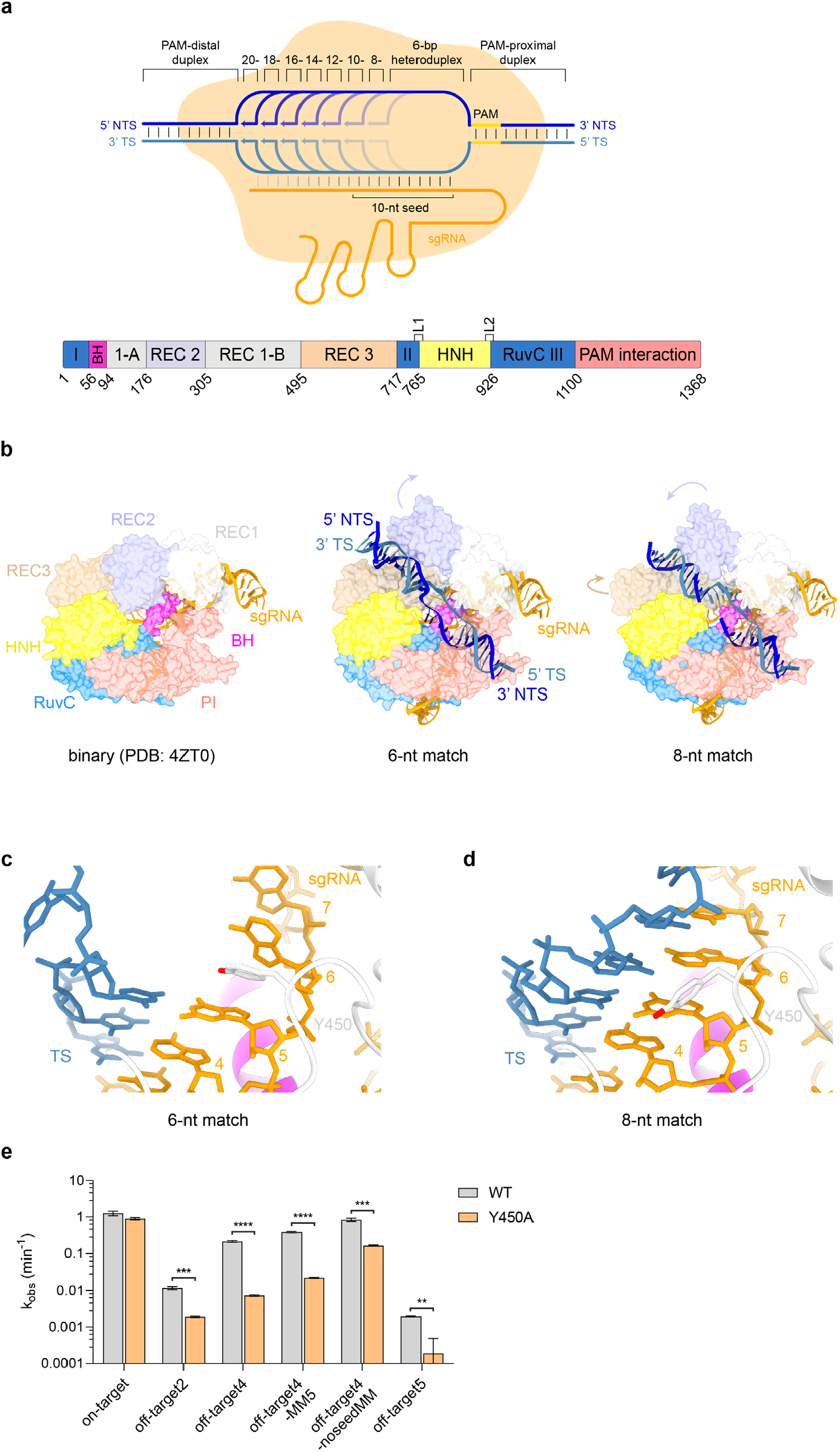
Target DNA binding induces restructuring of the Cas9 REC lobe. **a**, Top, Schematic depicting DNA-bound complexes with increasing complementarity to guide RNA. Bottom, Domain composition of the *Streptococcus pyogenes* Cas9 (SpCas9), BH: bridge helix. **b**, Structural comparison of the SpCas9 binary (left), 6-nt match (middle) and 8-nt match (right) complexes. **c**, Zoom-in view of the seed region of the guide RNA-target DNA heteroduplex in the 6-nt match complex. Tyr450 stacks between the 5^th^ and 6^th^ nucleotide, counting from the PAM-proximal end of the heteroduplex. **d**, Zoom-in view of the seed region of the guide RNA-target DNA heteroduplex in the 8-nt match complex. **e**, Cleavage rates of wild-type (WT) and Y450A mutant Cas9 against on- and off-target substrates. Data represents mean ± SEM (n=4), significance was determined by a two-tailed t-test, * p <0.05, ** = p <0.01, *** p <0.001, **** = p <0.0001.

Structural superpositions of the partially-matched complexes with the guide RNA-bound binary SpCas9 complex^12^ provide a framework for the visualization of the DNA binding mechanism, revealing stepwise domain rearrangements coupled to R-loop formation (**Extended Data Fig. 5a**). All complexes exhibit almost identical conformations of the bridge helix, REC1, RuvC, and PAM interaction domains, as well as the PAM-proximal dsDNA duplex and the sgRNA downstream (3’-terminal) of seed region. Conformational differences are observed in the positioning of the REC2, REC3, and the HNH domain relative to the emerging R-loop, consistent with the 3D variability analysis.

### R-loop initiation by bipartite seed sequence

The structure of the 6-nucleotide complementary target (6-nt match) complex reveals a 5-bp heteroduplex formed by the sgRNA seed sequence and TS DNA (**Fig. 1b**). Hybridization beyond the fifth seed sequence nucleotide is precluded by base stacking with the side chain of Tyr450, which was previously observed in the structure of the Cas9-sgRNA binary complex^12^ (**Fig. 1c**). Comparisons with the binary complex structure indicate that target strand hybridization is associated with the displacement of the REC2 domain out of the central binding channel (**Fig. 1b**). The PAM-distal duplex part of the DNA substrate is bound in a positively charged cleft formed by the REC2 and REC3 domains (**Fig. 1b, Extended Data Fig. 5b**), stabilized by interactions of the REC2 residues Ser219, Thr249 and Lys263 with the NTS backbone (**Extended Data Fig. 5c**), and REC3 residues Arg586 and Thr657 with the TS backbone (**Extended Data Fig. 5d**). Similar REC lobe conformation and protein contacts with the PAM-distal end of the DNA have been observed in a 3-bp heteroduplex complex described in a recent study^30^. Consequently, the NTS is positioned parallel to the guide RNA-TS DNA heteroduplex within the central binding channel (**Fig. 1b**). The 5’ terminal part of the sgRNA appears to be conformationally flexible but residual cryoEM density suggests its placement in a positively charged cleft located between the HNH and PAM-interaction domains (**Extended Data Fig. 5e**).

The structure of the 8-nucleotide complementary target (8-nt match) complex reveals that expansion of the R-loop heteroduplex, enabled by unstacking of Tyr450, forces further repositioning of the REC2 and REC3 domains to widen the binding channel as the PAM-distal duplex shifts deeper inside the channel (**Fig. 1d**; **Fig. 2a-c**; **Extended Data Fig. 5f**). R-loop propagation and PAM-distal duplex displacement results in the formation of new intermolecular contacts, with Cas9 contacting the PAM-distal duplex backbone through REC2 domain residues Ser217, Lys234 and Lys253, and REC3 residues Arg557 and Arg654 (**Extended Data Fig. 5g,h**).

**Figure 2.**
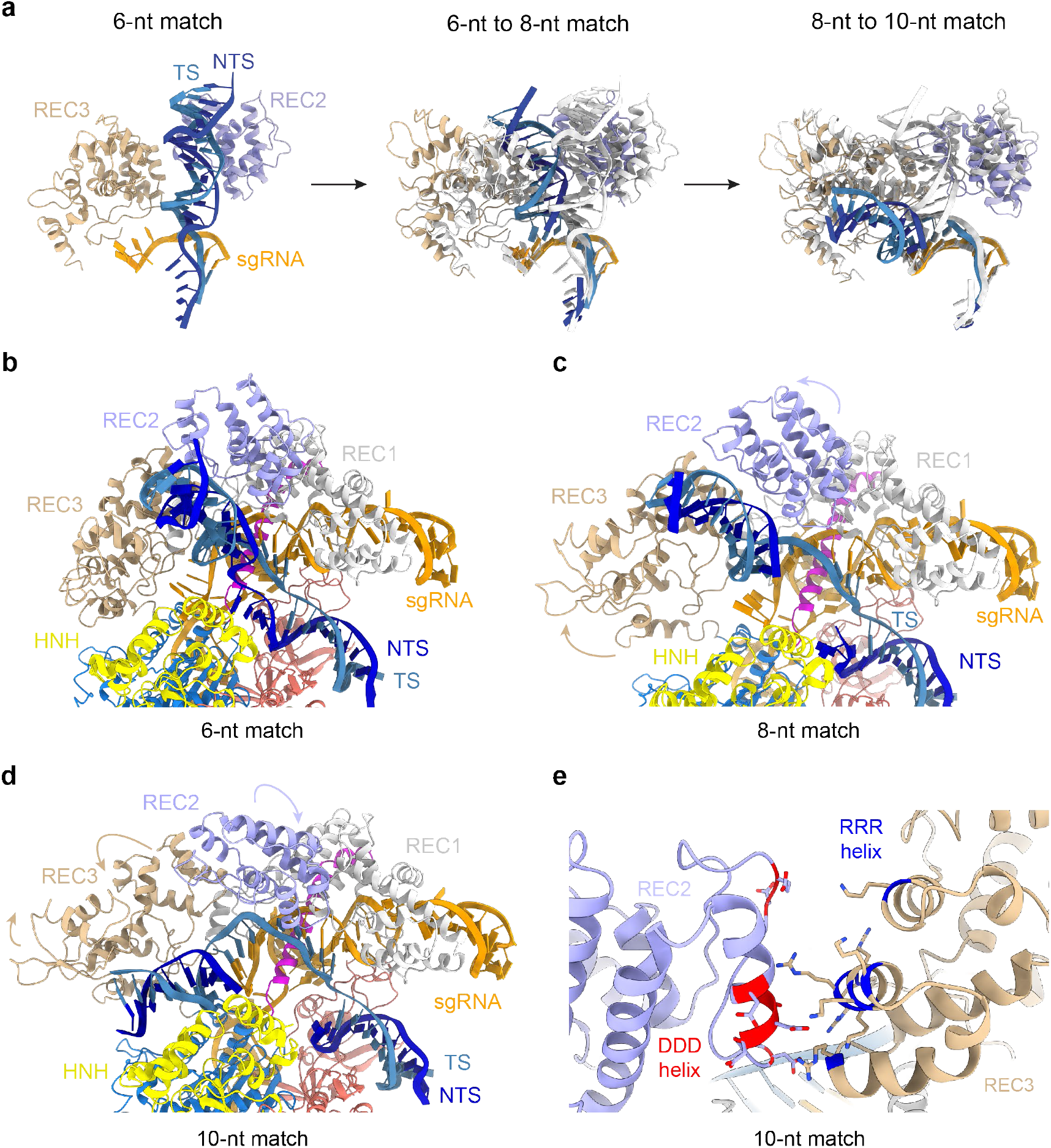
10-bp duplex formation drives DNA repositioning within Cas9. **a**, Zoom-in views of the conformational transitions in the PAM-distal DNA duplex and Cas9 REC2 and REC3 domains in the 6-, 8- and 10-nt match complex. **b**, Zoom-in view of the R-loop in the 6-nt match complex. **c**, Zoom-in view of the R-loop in the 8-nt match complex. **d**, Zoom-in view of the R-loop in the 10-nt match complex. **e**, Zoom-in view of the interaction between the REC2 domain DDD helix and the REC3 RRR helix.

Together, these observations suggest that the seed sequence of the Cas9 guide RNA is bipartite and that its hybridization with target DNA proceeds in two steps, consistent with the existence of a short-lived intermediate state observed in FRET studies^11,31^. To validate the observed interactions, we tested the cleavage activities of structure-based Cas9 mutant proteins in vitro (**Extended Data Fig. 6a**). Alanine substitution of Tyr450 resulted in significant reductions of off-target substrate cleavage, while on-target cleavage was unperturbed (**Fig. 1e**), confirming its role in seed sequence pre-ordering and suggesting that its perturbation leads to increased off-target discrimination. Interestingly, a subset of mutations of REC2 or REC3 residues resulted in increased off-target cleavage, as did the deletion of the REC2 domain (**Extended Data Fig. 6b**), in agreement with a previous study^31^. Collectively, these results underscore the importance of protein-DNA contacts during early steps of R-loop formation for the specificity of Cas9.

### R-loop propagation and remodelling

Further guide RNA-TS hybridisation to form a 10-bp heteroduplex causes a rearrangement of the REC2 and REC3 domains and repositioning of the PAM-distal DNA duplex into the positively charged central binding channel formed by the REC3, RuvC, and the HNH domains (**Fig. 2a**). Here, the PAM-distal dsDNA duplex forms a continuous base stack with the sgRNA-TS heteroduplex (**Fig. 2d**). The displaced NTS is positioned underneath the HNH domain and continues to run parallel to the extending guide RNA-TS DNA heteroduplex (**Extended Data Fig. 7a**). X-ray crystallographic analysis of the 10-nt match complex at a resolution of 2.8 Å (**Extended Data Table 2**) confirmed that the TS and NTS remain hybridised at the PAM-distal end of the DNA substrate (**Extended Data Fig. 7b**). The PAM-distal duplex is wedged between the REC3 and RuvC domains, and the L1 HNH linker (**Fig. 2d**; **Extended Data Fig. 7a-b**). The relocation of the PAM-distal duplex causes the REC2 domain to shift closer to the binding channel and occlude the cleavage site in TS DNA (**Fig. 2d**). This shift also establishes a new electrostatic interaction between a negatively charged helix in REC2 (Glu260, Asp261, Asp269, Asp272, Asp273, Asp274, Asp276) and a positively charged helix in REC3 (Lys599, Arg629, Lys646, Lys649, Lys652, Arg653, Arg654, Arg655), hereafter referred to as the DDD and RRR helices, respectively (**Fig. 2e**, which are highly conserved across Cas9 orthologs that contain a REC2 domain (**Extended Data Fig. 7c**) The off-target cleavage activity of Cas9 in vitro was significantly reduced by alanine substitutions of the interacting residues in the REC2 DDD helix, but not by mutations in the REC3 RRR helix (**Extended Data Fig. 6c**).

The HNH nuclease domain remains docked on the RuvC and PI domains in the 6-, 8- and 10-nt match complexes, with the active site buried at the interdomain interface (**Fig. 3a**). R-loop extension past the seed region to form a 12-bp heteroduplex does not result in major REC2/3 domain rearrangements, with the PAM-distal duplex remaining coaxially stacked onto the guide RNA-TS DNA heteroduplex throughout the 12-, 14-, 16- and 18-nt match complexes (**Fig. 3b**). In contrast, the HNH domain becomes disordered along with the surrounding RuvC 1011-1040 and PI 1245-1251 loops in the 12-nt match complex (**Fig. 3c**). Upon extension of the R-loop heteroduplex to 14 bp, the RuvC and PI loops responsible for HNH docking remain structurally disordered (**Fig. 3c, Extended Data Fig. 8a**) and residual density is observed for the HNH domain as its L2 linker contacts the guide RNA-TS heteroduplex (**Extended Data Fig. 8a**). Further extension of the R-loop heteroduplex from 14- to 16-bp causes translocation of the HNH domain towards the guide RNA-TS DNA heteroduplex within the central binding channel (**Fig. 3c**). Facilitated by the formation of the PAM-distal part of the R-loop, a loop in the RuvC domain (residues 1030-1040) restructures into a helical conformation, establishing interactions with the L2 linker (**Extended Data Fig. 8b**). This repositions the L2 linker and shifts the HNH domain on top of the heteroduplex, sealing off the central binding channel (**Fig. 3c, Extended Data Fig. 8c**). The HNH domain remains in a catalytically incompetent orientation, with its active site located ∼31 Å away from the scissile phosphate group in the TS.

**Figure 3.**
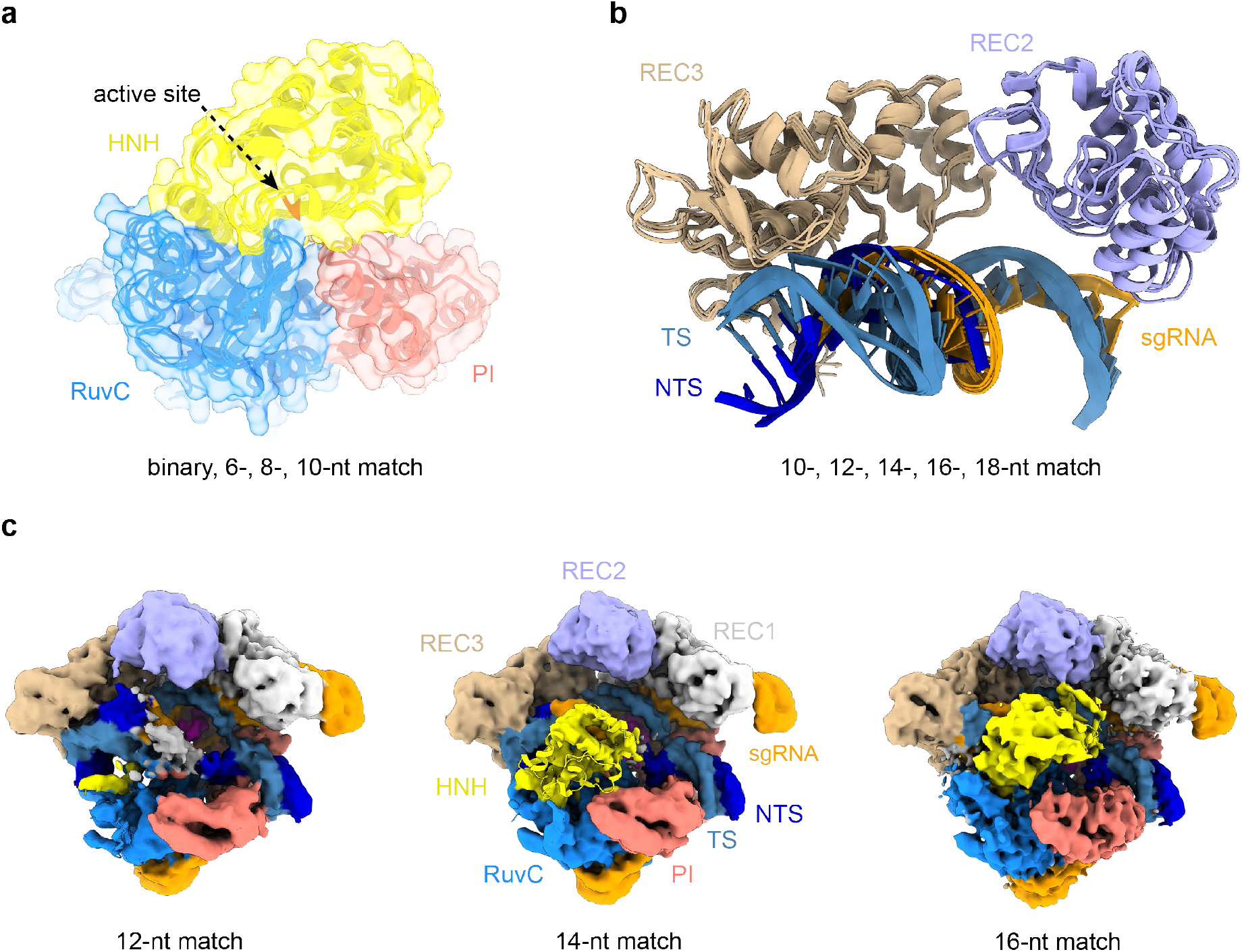
Target pairing past the seed region undocks the HNH nuclease domain. **a**, Position of the HNH catalytic site in the binary, 6-, 8-, and 10-nt match complexes. **b**, Structural overlay of the REC2 and REC3 domains in the 10-, 12-, 14-, 16- and 18-nt match (checkpoint) complexes. **c**, Overview of the 12-nt match (left), 14-nt match (middle) and 16-nt match (right) complexes, shown in the same orientations. For each complex, the unsharpened cryo-EM map is overlaid with the respective atomic model. The 12-nt match complex map shows residual density for the displaced NTS (white); The 14-nt match map reveals residual density corresponding to the HNH domain; no density is visible for NTS. Cryo-EM maps are coloured according the schematic in **Fig. 1a**.

### Conformational checkpoint and nuclease activation

Previous studies have shown that substrates containing 4-bp mismatches at the PAM-distal end of the target sequence (positions 16-20) are generally refractory to Cas9 cleavage, whereas substrates containing mismatches at positions 19-20 are efficiently cleaved^13,23,24,32^. The cryo-EM reconstruction of the 18-nt match complex in the presence of 1 mM Mg^2+^ reveals that the most populated 3D class in the sample represents a pre-cleavage state with an intact TS and disordered NTS (**Fig. 4a**). Upon extension of the R-loop to 18 bp, the HNH domain continues to assume the catalytically incompetent orientation observed in the 16-nt match complex, while the conformation of the REC2/3 domains remains the same as in the 12-, 14- and 16-nt match complexes (**Fig. 3b, 4a**). The observed conformation is thus consistent with a catalytically inactive checkpoint state inferred from prior biophysical and structural studies^23,24,32^.

**Figure 4.**
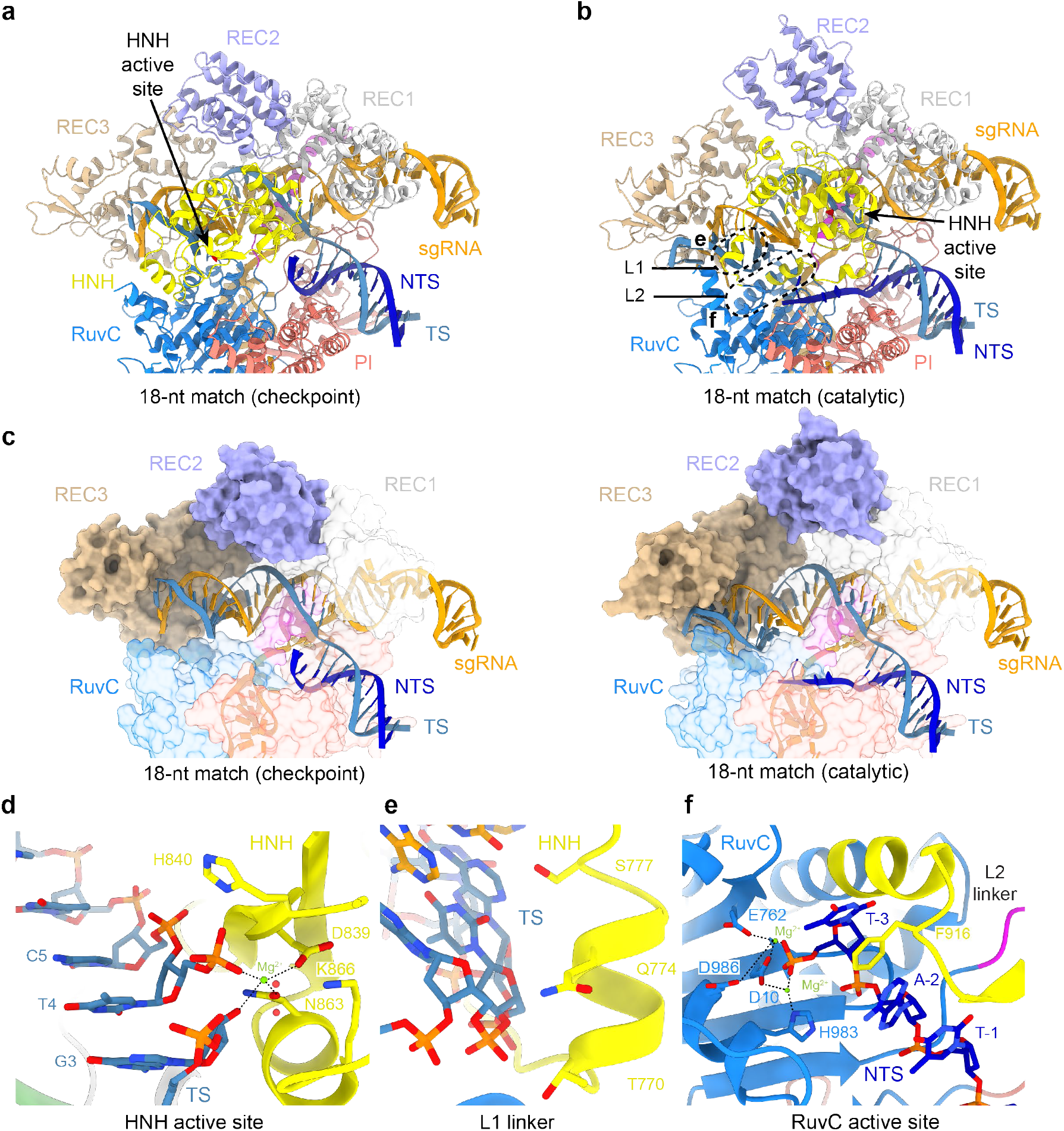
HNH rotation is required TS cleavage and RuvC nuclease activation. **a**, Structure of the 18-nt match complex in the pre-cleavage, checkpoint state. **b**. Structure of the 18-nt match complex in the catalytically active state. **c**, Conformations of the guide-target heteroduplexes and REC2/REC3 domains in the 18-nt match checkpoint (left) and catalytic (right) complexes. The structures are shown in the same orientations as in panels a,b. The HNH domain has been omitted from the images for clarity. **d**, Zoom-in view of the HNH nuclease active site in the 18-match catalytic complex containing bound cleaved target strand (TS). **e**, Zoom-in view of the L1 linker contacting the minor groove of the guide RNA-target DNA heteroduplex. **f**, Zoom-in view of the RuvC nuclease active site containing the 3’-terminal product of cleaved non-target strand (NTS).

The cryo-EM reconstruction obtained from a sample reconstituted in the presence of 10 mM Mg^2+^ reveals a catalytically active conformation in which both the TS and the NTS are cleaved at the expected positions (**Fig. 4b-f**). In contrast to previously reported structures of catalytically active Cas9 enzymes^28,33,34^, the PAM-proximal part of the cleaved NTS remains bound in the RuvC active site (**Fig. 4b,f**). In this state, the REC2 domain is shifted away from the TS cleavage site, enabling the HNH domain to undergo a ∼140° rotation to engage the TS scissile phosphate with its active site and catalyse its hydrolysis via a one-metal-ion mechanism (**Fig. 4d**), in agreement with prior structural data^28^. This rearrangement is facilitated by pronounced bending of the PAM-distal region of the guide RNA-TS DNA heteroduplex and a concomitant reorientation of the REC3 domain that preserves interactions with the heteroduplex (**Fig 4c**). HNH domain rotation is brought about by restructuring of the L1 and L2 linkers, which results in the widening of the NTS binding cleft and exposure of the RuvC active site (**Fig. 4b,e,f**). The L1 linker, which is structurally disordered in the 18-nt match checkpoint complex, forms an α-helix and interacts with the minor groove of the guide RNA-TS DNA heteroduplex via multiple hydrogen bonding interactions (**Fig. 4e**). The L2 linker helix becomes extended, allowing Phe916 to intercalate between NTS nucleobases by π-π stacking, thereby stabilizing the NTS in the RuvC active site (**Fig. 4f**). The NTS scissile phosphate is coordinated by two Mg^2+^ ions, its position consistent with a His983-dependent catalytic mechanism proposed by molecular dynamics simulations^35^. Together, these structural observations provide a rationale for the allosteric coupling of R-loop formation with HNH domain rearrangement and RuvC active site accessibility, in agreement with single-molecule studies showing that PAM-distal end positioning modulates HNH domain conformation^32^. The concerted mechanism of HNH domain and NTS repositioning has also been reported in a complementary structural study^36^.

## Conclusions

In sum, our structural analysis of SpCas9 along its DNA binding pathway points to a mechanism whereby R-loop formation is allosterically and energetically coupled to domain rearrangements necessary for nuclease domain activation (**Extended Data Fig. 9**). The initial phase of R-loop formation is facilitated by target strand hybridization to a bipartite seed sequence of the guide RNA and interactions of the PAM-distal DNA with the Cas9 REC2 and REC3 domains. The observation of a bipartite seed sequence in the Cas9 guide RNA and a two-step seed hybridization mechanism involving a conformational rearrangement brings parallels with other RNA-guided nucleic acid-targeting systems including the Cascade complex and Argonaute proteins, both of which feature discontinuous seed sequences in their guide RNAs^37–41^. We identify mutations that destabilise the binding intermediate states and thus increase off-target discrimination, which presents an opportunity for the development of novel high-fidelity SpCas9 variants. As most off-target sequences are only bound but not cleaved^19–21,42^, these variants could prove useful for applications that rely on the fidelity of Cas9 target binding^43–46^, such as transcriptional regulation or base editing^47^. Directional target DNA hybridization is associated with dynamic repositioning of the REC2/3 and HNH domains to initially assume a catalytically inactive, checkpoint conformation upon R-loop completion. As conformational activation of the nuclease domains is allosterically controlled by structural distortion of the PAM-distal end of the guide-target heteroduplex and the sensing of its integrity by Cas9, it is precluded by incomplete PAM-distal heteroduplex pairing (<18 bp). Bona fide off-target substrates are able to pass the conformational checkpoint because they maintain heteroduplex integrity despite the presence of PAM-distal mismatches, in agreement with our recent structural data^48^. Furthermore, guide RNA modifications that result in altered heteroduplex conformation have profound effects on Cas9 nuclease activity and specificity^49^. Together, our structural studies thus highlight the importance of maintaining guide-target complementarity and proper heteroduplex geometry, consistent with biophysical and computational studies showing that the conformation of the R-loop heteroduplex strongly affects off-target binding^11,50^. These findings thus have important implications for ongoing experimental and computational studies of CRISPR-Cas9 off-target activity, and will inform its further technological development.

## Methods

### Expression and purification of Cas9 proteins

WT and mutant *Streptococcus pyogenes* Cas9 proteins were expressed in *E*.*coli* Rosetta 2 (DE3) (Novagen) for 16 hours at 18 °C as fusion proteins with an N-terminal His_6_-MBP-TEV tag. Bacterial pellets were resuspended and lysed in 20 mM HEPES-KOH pH 7.5, 500 mM KCl, 5 mM imidazole, and protease inhibitors. Cell lysates were clarified using ultracentrifugation and loaded on a 15 ml Ni-NTA Superflow column (QIAGEN) and washed with 7 column volumes of 20 mM HEPES-KOH pH 7.5, 500 mM KCl, 5 mM imidazole. Tagged Cas9 was eluted with 10 column volumes of 20 mM HEPES-KOH pH 7.5, 250 mM KCl, 200 mM imidazole. Salt concentration was adjusted to 250 mM KCl and the protein was loaded on a 10 ml HiTrap Heparin HP column (GE Healthcare) equilibrated in 20 mM HEPES-KOH pH 7.5, 250 mM KCl, 1 mM DTT. The column was washed with 5 column volumes of 20 mM HEPES-KOH pH 7.5, 250 mM KCl, 1 mM DTT, and dCas9 was eluted with 15 column volumes of 20 mM HEPES-KOH pH 7.5, 1.5 M KCl, 1 mM DTT, in a 0-50% gradient (peak elution around 500 mM KCl). His6-MBP tag was removed by TEV protease cleavage overnight at 4 °C with gentle shaking. The untagged protein was concentrated and further purified on a Superdex 200 16/600 gel filtration column (GE Healthcare) in 20 mM HEPES-KOH pH 7.5, 500 mM KCl, 1 mM DTT. Pure fractions were concentrated to 10 mg/ml, flash frozen in liquid nitrogen and stored at 80 °C.

### sgRNA *in vitro* transcription

The single guide RNA (sgRNA) was transcribed from a dsDNA template in a 5 ml transcription reaction (30 mM Tris-HCl pH 8.1, 25 mM MgCl2, 2 mM spermidine, 0.01% Triton X-100, 5 mM CTP, 5 mM ATP, 5 mM GTP, 5 mM UTP, 10 mM DTT, 1 μM DNA transcription template, 0.5 units inorganic pyrophosphatase (Thermo Fisher), 250 μg T7 RNA polymerase). The transcription reaction was incubated at 37 °C for 5 hours, after which the dsDNA template was degraded for 30 minutes with 15 units of RQ1 DNAse (Promega). The transcribed sgRNA was PAGE purified on an 8% denaturing polyacrylamide gel containing 7 M urea, ethanol precipitated and dissolved in DEPC-treated water.

### Gel filtration binding assay

The dCas9-gRNA complex was assembled by incubating 371 picomoles of dCas9 with 400 picomoles of the sgRNA in 20 mM HEPES-KOH pH 7.5, 200 mM KCl, 2 mM MgCl_2_ for 10 minutes at room temperature. Then 250 picomoles of Cy5-labeled dsDNA substrate was added and incubated another 15 minutes. The volume was adjusted up to 100 μl with reaction buffer and the mixture was centrifuged to remove possible precipitates. Individual reactions were transferred to a 96-well plate and analysed using a Superdex 200 Increase 5/150 GL gel filtration column (GE Healthcare) attached to an Agilent 1200 Series Gradient HPLC system. The 260 nm, 280 nm, and Cy5 signals were exported and plotted as a function of the retention volume in GraphPad Prism 9.

### In vitro nuclease activity assays

Cleavage reactions were performed at 37 °C in reaction buffer, containing 20 mM HEPES pH 7.5, 250 mM KCl, 5 mM MgCl_2_ and 1 mM DTT. First, Cas9 protein was pre-incubated with sgRNA in 1:1.25 ratio for 10 minutes at room temperature. The protein-RNA complex was rapidly mixed with the ATTO-532 labelled dsDNA, to yield final concentrations of 1.67 μM protein and 66.67 nM substrate in a 7.5 μl reaction. Time points were harvested at 1, 2.5, 5, 15, 45, 90, 150 minutes, and 24 hours. Cleavage was stopped by addition of 2 μl of 250 mM EDTA, 0.5% SDS and 20 μg of Proteinase K. Formamide was added to the reactions with final concentration of 50%, samples were incubated at 95 °C for 10 minutes, and resolved on a 15% denaturing PAGE gel containing 7M urea and imaged using a Typhoon FLA 9500 gel imager. Rate constants (k_obs_) were extracted from single exponential fits: [Product] = A*(1-exp(-k_obs_*t)). Data are presented as mean ± s.e.m. (*n*=4 biologically independent replicates). Statistical analysis was performed using a two-sided t test. The interval of confidence used was 95%.

### Crystallisation and X-ray structure determination

The 10-nt complementary ternary complex of dCas9 was assembled by first incubating dCas9 with the sgRNA in a 1:1.5 molar ratio, and pre-purifying the binary complex on a Superdex 200 16/600 gel filtration column (GE Healthcare) in 20 mM HEPES-KOH pH 7.5, 500 mM KCl, 1 mM DTT. The binary complex was diluted in 20 mM HEPES-KOH pH 7.5, 250 mM KCl, 1 mM DTT to 2.5 mg/ml and the partially complementary dsDNA substrate was added in 1:1.5 molar excess. For crystallisation, 1 μl of the ternary complex (1.5-2.5 mg/ml) was mixed with 1 μl of the reservoir solution (0.1 M sodium cacodylate pH 6.5, 0.8-1.2 M ammonium formate, 12-14% PEG4000) and crystals were grown at 20 °C using the hanging drop vapour diffusion setup. Crystals were harvested after 3-4 weeks, cryoprotected in 0.1 M Na cacodylate pH 6.5, 1.0 M ammonium formate, 13% PEG4000, 20% glycerol, 2 mM MgCl_2_, and flash-cooled in liquid nitrogen. Diffraction data was measured at the beamline PXIII of the Swiss Light Source at a temperature of 100 K (Paul Scherrer Institute, Villigen, Switzerland) and processed using the autoPROC and STARANISO package with anisotropic cut-off^51^. Phases were obtained by molecular replacement using the Phaser module of the Phenix package^52^ using the NUC lobe of the PDB ID: 5FQ5 as initial search model. The crystals belonged to the P1 space group and contained two copies of the complex in the asymmetric unit.

### Cryo-EM sample preparation and data acquisition

To assemble the 6-, 8-, 10-, 12-, 14-, and 16-nt match complexes, dCas9 protein was mixed with the sgRNA in a 1:1.5 molar ratio, and incubated at room temperature for 10 minutes in buffer 20 mM HEPES-KOH pH 7.5, 250 mM KCl, 1 mM DTT. The respective partially complementary dsDNA substrate was then added in a 1:3 Cas9:DNA molar ratio and incubated another 20 minutes at room temperature. The complexes were then purified using a Superdex 200 Increase 10/300 GL gel filtration column (GE Healthcare) and eluted in 20 mM HEPES-KOH pH 7.5, 250 mM KCl, 1 mM DTT. Concentration of the monomeric peak was determined using the Qubit 4 Fluorometer Protein Assay, and then diluted to 0.275 mg/ml in 20 mM HEPES-KOH pH 7.5, 250 mM KCl cold buffer. 3 μl of diluted complexes were applied to a glow discharged 200-mesh holey carbon grid (Au 1.2/1.3 Quantifoil Micro Tools), blotted for 1.5-2.5 s at 90% humidity, 20 °C, plunge frozen in liquid propane/ethane mix (Vitrobot, FEI) and stored in liquid nitrogen. To prepare the 18-nt match (checkpoint), wild-type Cas9-sgRNA complex was reconstituted with substrate DNA in 20 mM HEPES-KOH pH 7.5, 150 mM KCl, 1 mM DTT buffer, and incubated with 1 mM MgCl_2_ for 1 minute at 37 °C prior to vitrification. The 18-nt match catalytic complex was reconstituted in 20 mM HEPES-KOH pH 7.5, 100 mM KCl, 1 mM DTT buffer, and incubated with 10 mM MgCl_2_ for 1 minute at 37 °C prior to vitrification. Data collection was performed on a 300 kV FEI Titan Krios G3i microscope equipped with a Gatan Quantum Energy Filter and a K3 direct detection camera in super-resolution mode. Micrographs were recorded at a calibrated magnification of 130,000 x with a pixel size of 0.325 Å and subsequently binned to 0.65 Å. Data acquisition was performed automatically using EPU with three shots per hole at −0.8 μm to −2.2 μm defocus. Data for the 18-nt match (checkpoint) complex was collected using a Titan Krios G4 equipped with a SelectrisX energy filter and a FalconIV detector at a magnification of 270,000 x, pixel size of 0.45 Å, defocus −0.8 μm to −1.5 μm.

### Cryo-EM data processing

Acquired super-resolution cryoEM data was processed using cryoSPARC^53^. Gain-corrected micrographs were imported and binned to a pixel size of 0.65 Å during patch motion correction. After patch CTF estimation, micrographs with a resolution estimation worse than 5 Å and full-frame motion distance larger than 100 Å were discarded. Initial particles were picked using blob picker with 100-140 Å particle size. Particle picks were inspected and particles with NCC scores below 0.4 were discarded. Remaining particles were extracted with a box size of 384 × 384 pixels, down-sampled to 192 × 192 pixels. After 2D classification, templates were generated using good classes and particle picking was repeated using the template picker. Duplicate particles were removed, and 2D classified Cas9 particles were used for *ab initio* 3D reconstruction. All partially bound complexes displayed several conformational states. After several rounds of 3D classification, classes with most detailed features were reextracted using full 384 × 384 pixel box size and subjected to non-uniform refinement to generate high-resolution reconstructions^54^. The 18-nt match (checkpoint) complex was extracted with a box size of 504 × 504. Each map was sharpened using the appropriate B-factor value to enhance structural features, and local resolution was calculated and visualised using ChimeraX^55^.

### Structural model building, refinement, and analysis

Manual Cas9 domain placement, model adjustment and nucleic acid building was completed using COOT^56^. Atomic model refinement was performed using Phenix.refine for X-ray data^57^ and Phenix.real_space_refine for cryoEM^58^. The quality of refined models was assessed using MolProbity^59^. Protein-nucleic acid interactions were analysed using the PISA web server^60^. Characterisation of the guide-protospacer duplex was performed using the 3DNA 2.0 web server^61^. Structural figures were generated using ChimeraX^55^.

### Protein sequence alignment

Protein sequences of Cas9 orthologues harbouring the REC2 domain were obtained from UniProt^62^. Sequence alignment was performed using MUSCLE with default parameters^63^. Alignment was visualised using Jalview with highlighting only the conservation of charged residues^64^.

## Data Availability

Atomic coordinates, maps and structure factors of the reported X-ray and cryo-EM structures have been deposited in the PDB under accession numbers 7Z4D (10-nt match complex, X-ray), 7Z4C (6-nt match complex, cryo-EM), 7Z4E (8-nt match complex, cryo-EM), 7Z4K (10-nt match complex, cryo-EM), 7Z4G (12-nt match complex, cryo-EM), 7Z4H (14-nt match complex, cryo-EM), and 7Z4I (16-nt match complex, cryo-EM), 7Z4L (18-nt match checkpoint complex, cryo-EM), and 7Z4J (18-nt match catalytic complex, cryo-EM) and in the EMDB under accession codes 14493 (6-nt match complex, cryo-EM), 14494 (8-nt match complex, cryo-EM), 14500 (10-nt match complex, cryo-EM), 14496 (12-nt match complex, cryo-EM), 14497 (14-nt match complex, cryo-EM), 14498 (16-nt match complex, cryo-EM), 14501 (18-nt match checkpoint complex, cryo-EM), and 14499 (18-nt match catalytic complex, cryo-EM).

## Author contributions

M.P. and M.J. conceived the study and designed experiments. M.P. purified Cas9, performed *in vitro* cleavage assays, crystallized 10-bp heteroduplex complex, prepared cryo-EM samples and solved the structures. I.Q., L.L. and L.M.M. expressed and purified mutant constructs, assisted with figure preparation, and collected cryo-EM data. M.S. collected cryo-EM data. M.P. and M.J. performed structural analysis and wrote the manuscript, with input from the remaining authors.

## Acknowledgements

This work was supported by the Swiss National Science Foundation Grant 31003A_182567 (to M.J.). M.J. is an International Research Scholar of the Howard Hughes Medical Institute and Vallee Scholar of the Bert L & N Kuggie Vallee Foundation. We thank Simona Sorrentino (UZH Center for Microscopy and Image Analysis) and Alexander Myasnikov (Dubochet Center for Imaging, Lausanne) for their assistance with cryogenic electron microscopy data collection. We thank Franziska Boneberg and Christelle Chanez for their help with preparing reagents. We thank members of the Jinek laboratory for discussion and critical reading of the manuscript. We are grateful to Josh Cofsky, Katarzyna Soczek and Jennifer Doudna for sharing unpublished data and helpful comments.

---

**Extended Data Figure 1.**
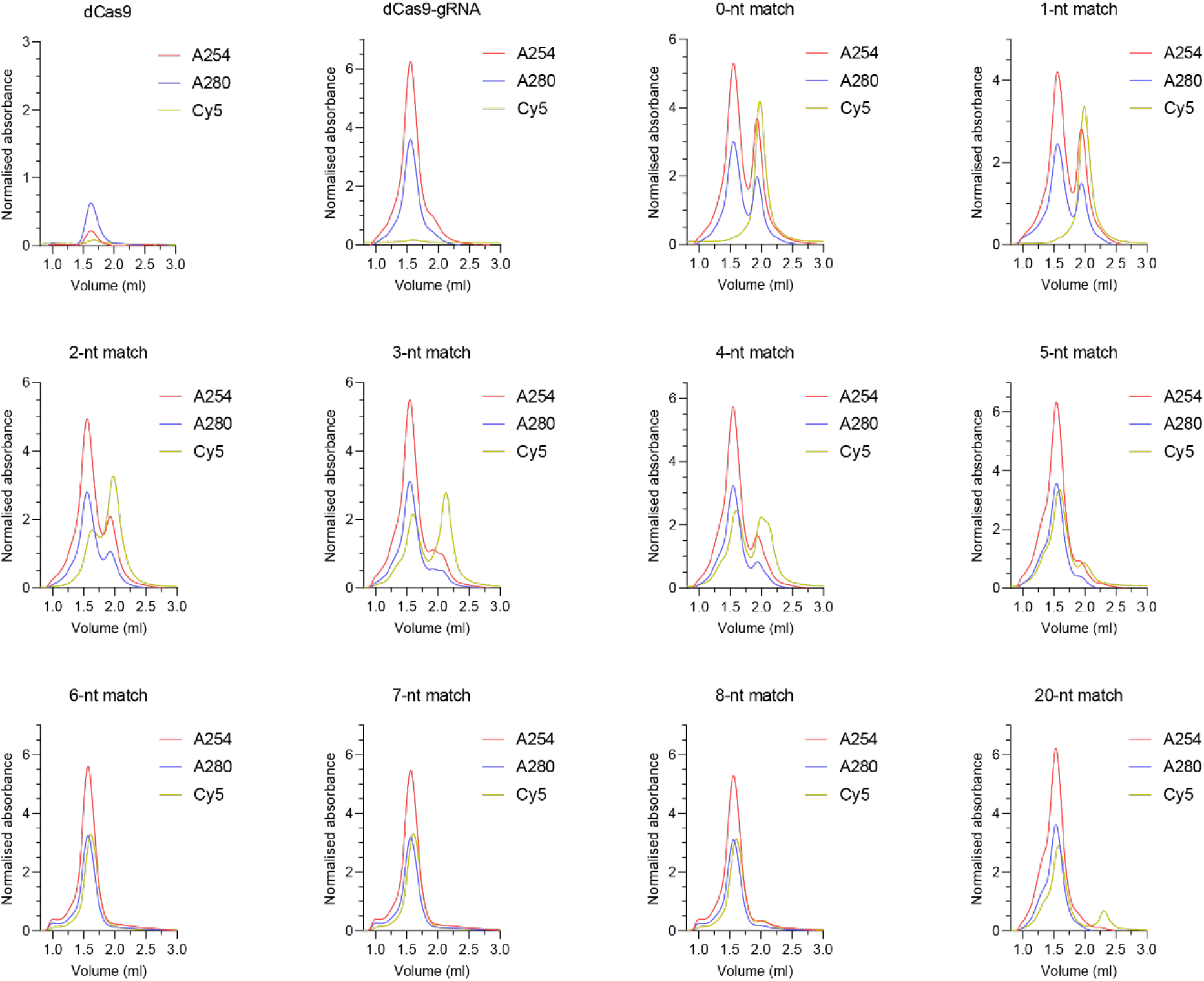
Minimal target complementarity necessary for stable Cas9 binding. Size exclusion chromatography analysis of nuclease-dead SpCas9 complexed with sgRNA and Cy5-labeled DNA substrates with increasing extent of guide-target complementarity. A254/A280 and Cy5 signals were normalised.

**Extended Data Figure 2.**
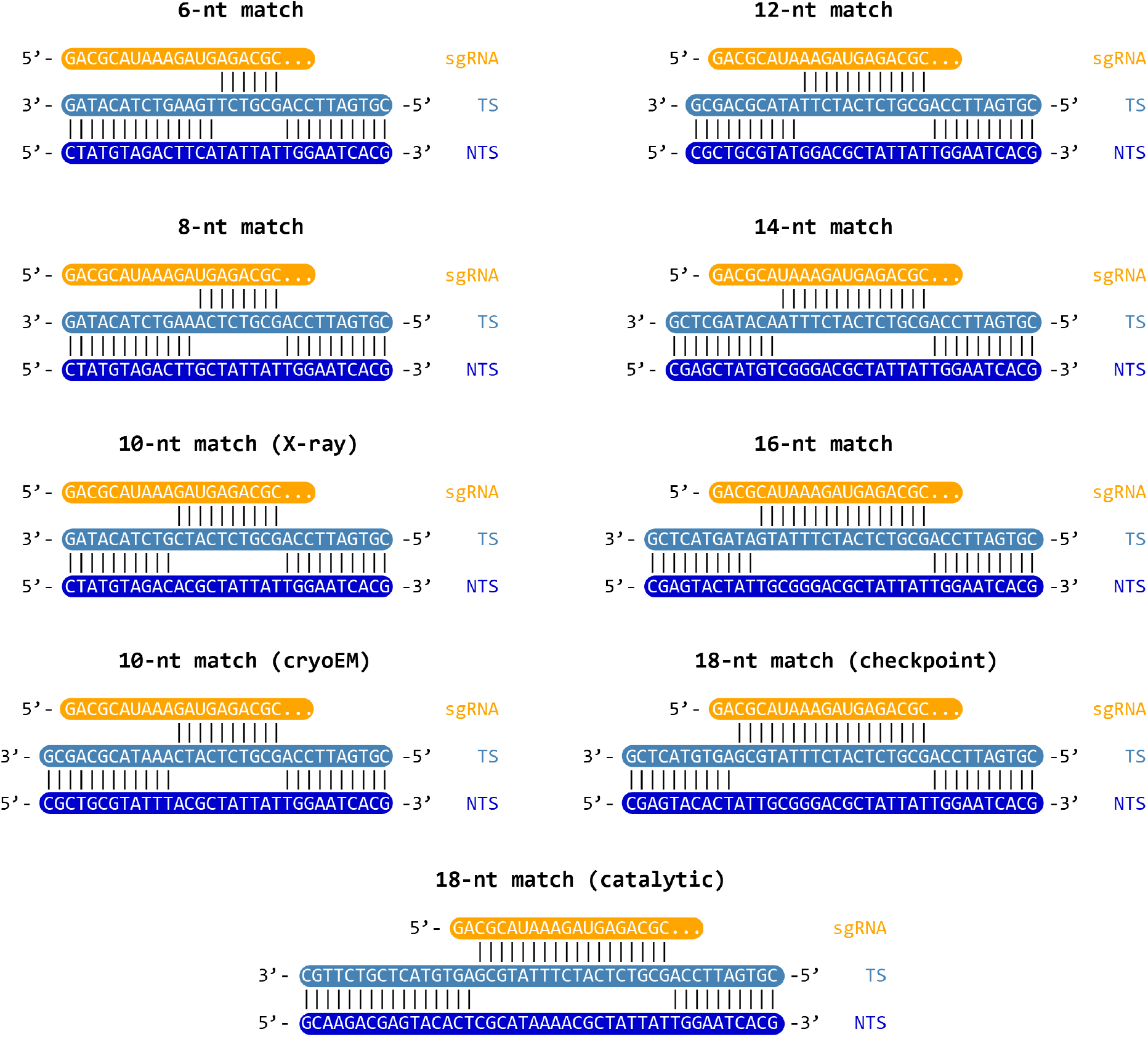
Schematic representation of DNA substrates used in structural studies. Base pair complementarity between sgRNA, target strand (TS), and non-target strand (NTS) is indicated by black lines.

**Extended Data Figure 3.**
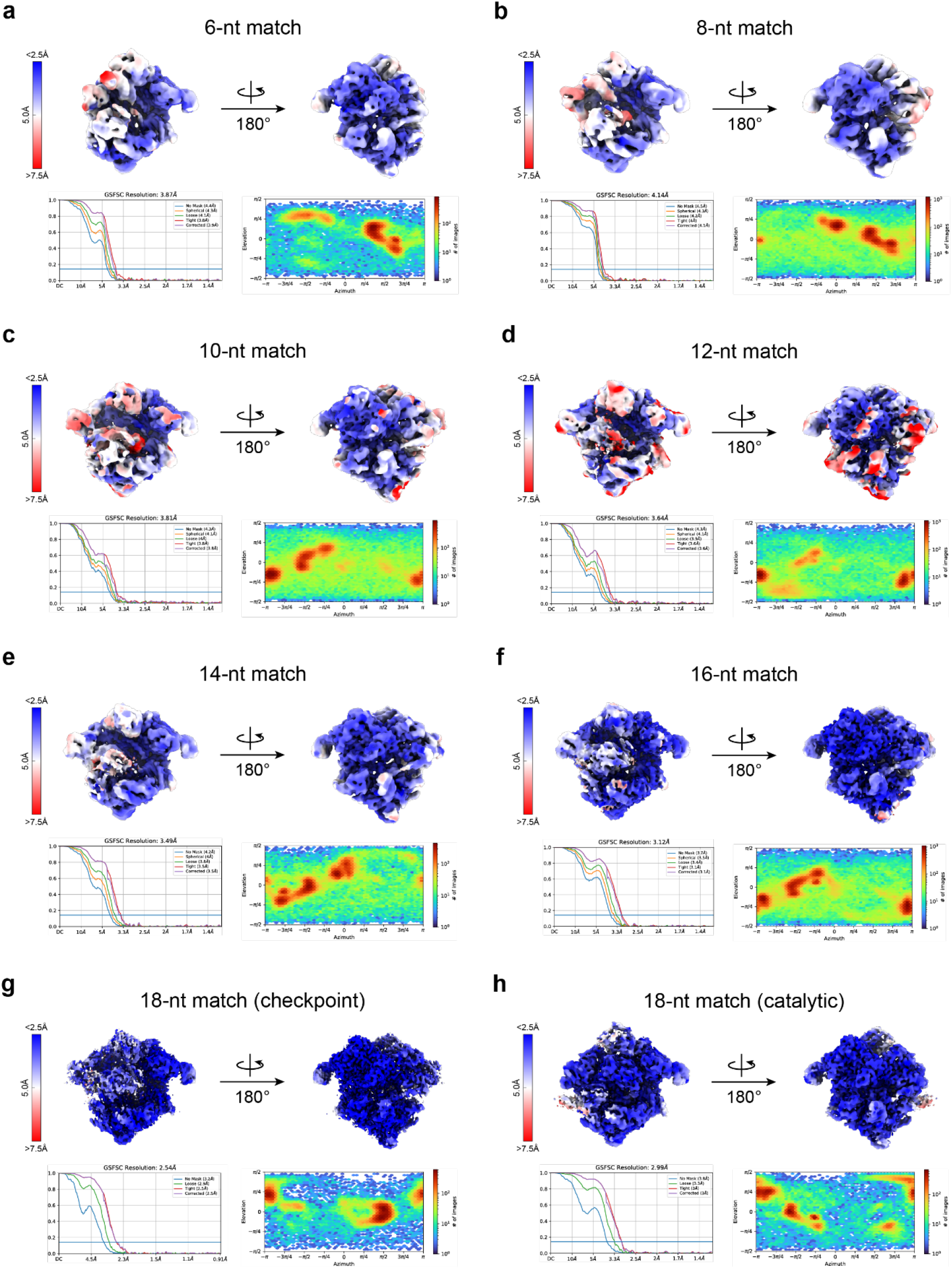
Cryo-EM density maps of partial R-loop SpCas9 complexes. Front (left) and back (right) views of unsharpened cryo-EM density maps of the partially-bound Cas9 complexes. Maps are coloured by local resolution, and gold-standard FSC of 0.143 resolution graphs and particle distribution heatmaps are indicated for each complex.

**Extended Data Figure 4.**
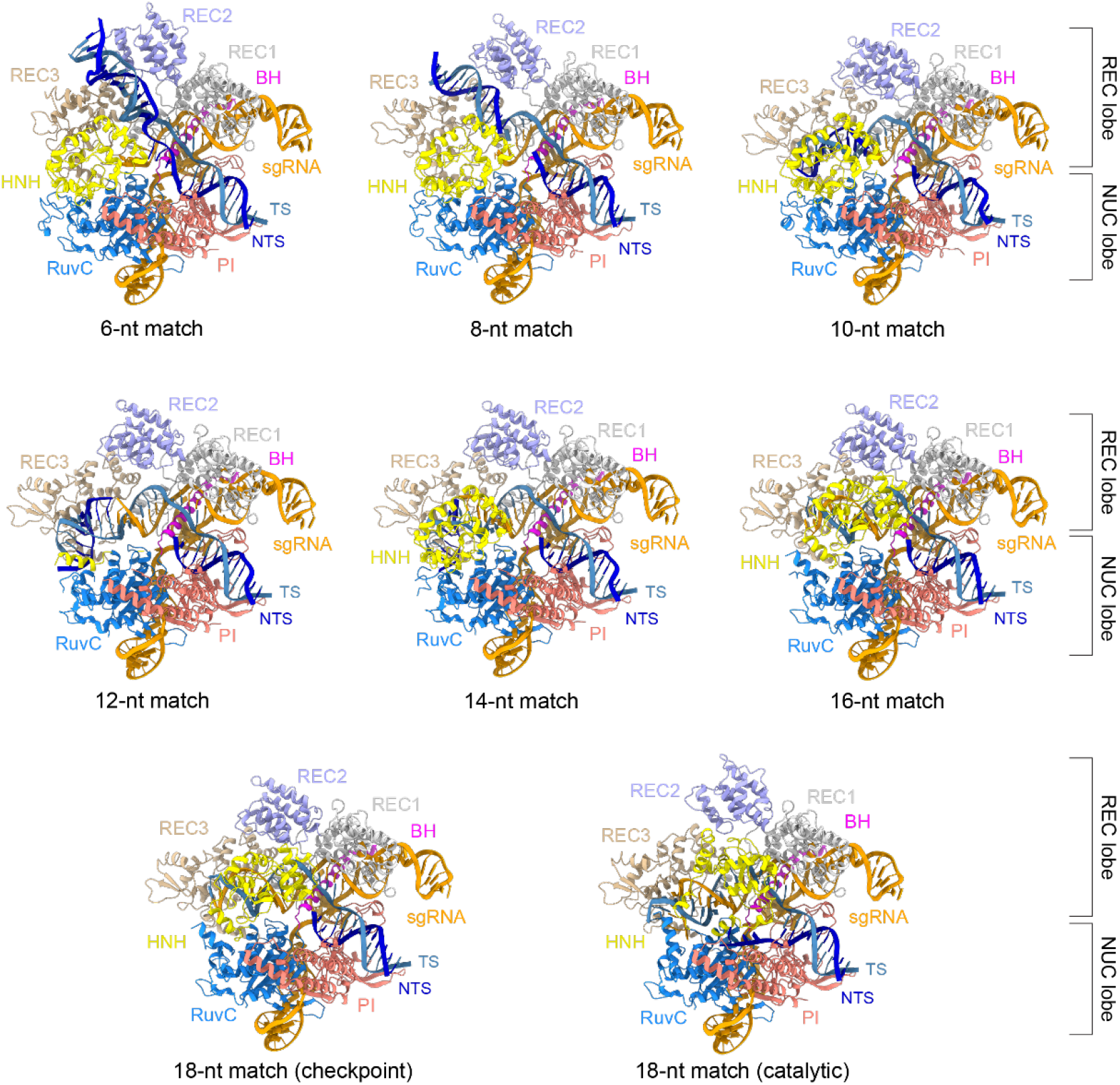
Structural models of SpCas9 partially-bound target complexes. Cartoon representations of target DNA-bound 6-, 8-, 10-, 12-, 14-, 16-, 18-nt match (checkpoint and catalytic) complexes of SpCas9. Each model was generated based on the corresponding map shown in **Extended Data Figure 3**.

**Extended Data Figure 5.**
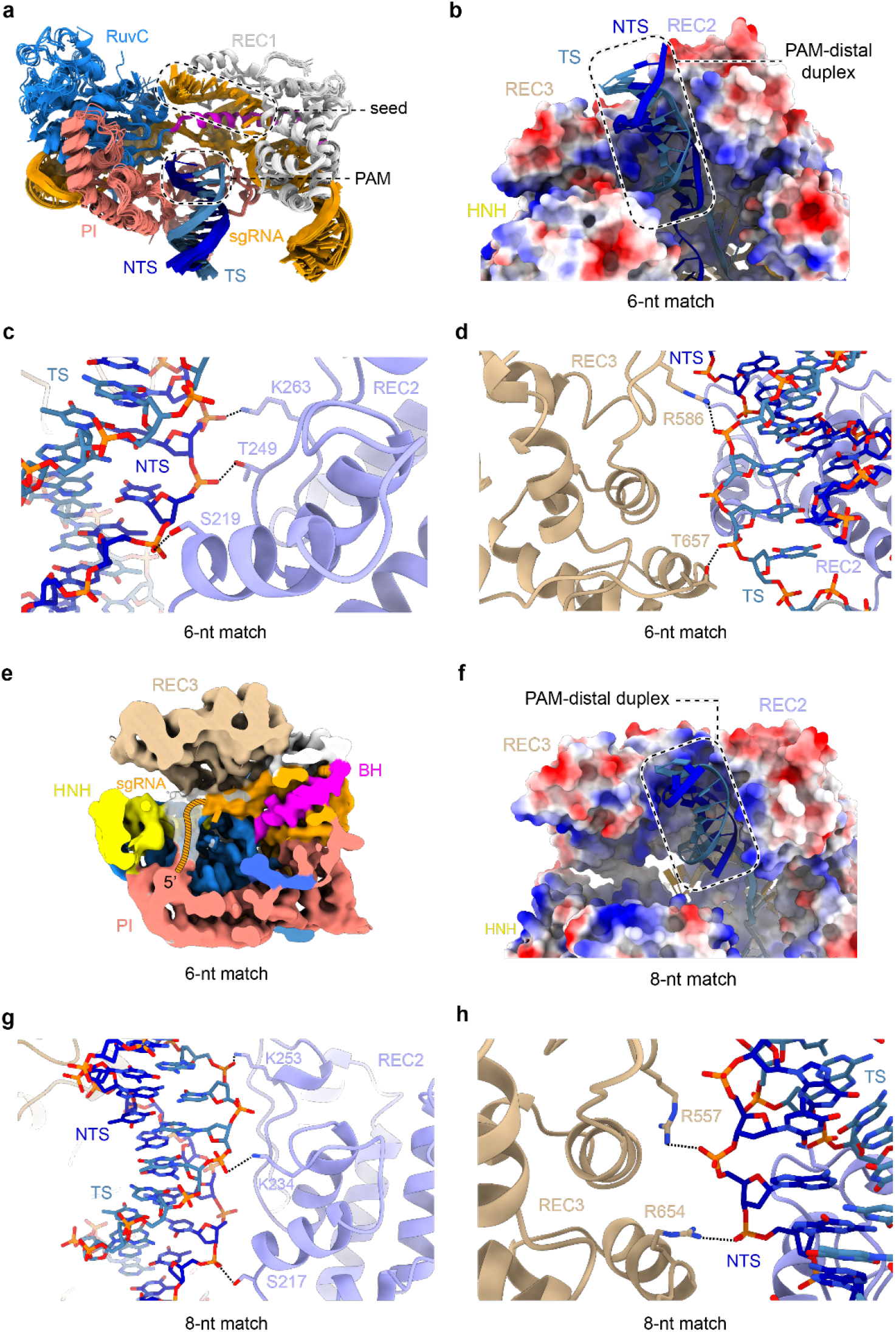
Stabilisation of the PAM-distal duplex by REC2/3 domains. **a**, Structural overlays of the SpCas9 bridge helix (BH), REC1, RuvC, and PI domains, as well as the PAM-proximal DNA duplex and the sgRNA from the partially bound complex structures determined by crystallography and cryoEM, and full R-loop complexes (PDB: 6O0X, 6O0Y, 6O0Z) (Zhu et al., 2019a). **b**, Zoom-in view of the PAM-distal DNA duplex in the 6-nt match complex. The protein surface is coloured according to electrostatic surface potential, with red denoting negative and blue positive charge. **c**, Interactions between SpCas9 REC2 domain and the backbone of the PAM-distal NTS in the 6-nt match complex. **d**, Interactions between the REC3 domain and the backbone of the PAM-distal TS in the 6-nt match complex. **e**, Central slice through the 6-nt match complex. Cryo-EM density map is coloured according to **Fig. 1a**. White density indicates positioning of the 5’ sgRNA end. **f**, PAM-distal DNA duplex in the 8-nt match complex remains positioned in a positively charged cavity between the REC2 and REC3 domains. The protein surface is coloured according to electrostatic surface potential. **g**, Interactions of the REC2 domain with the PAM-distal DNA duplex in the 8-nt match complex. **h**, Interactions of the REC3 domain with the NTS of the PAM-distal duplex in the 8-nt match complex.

**Extended Data Figure 6.**
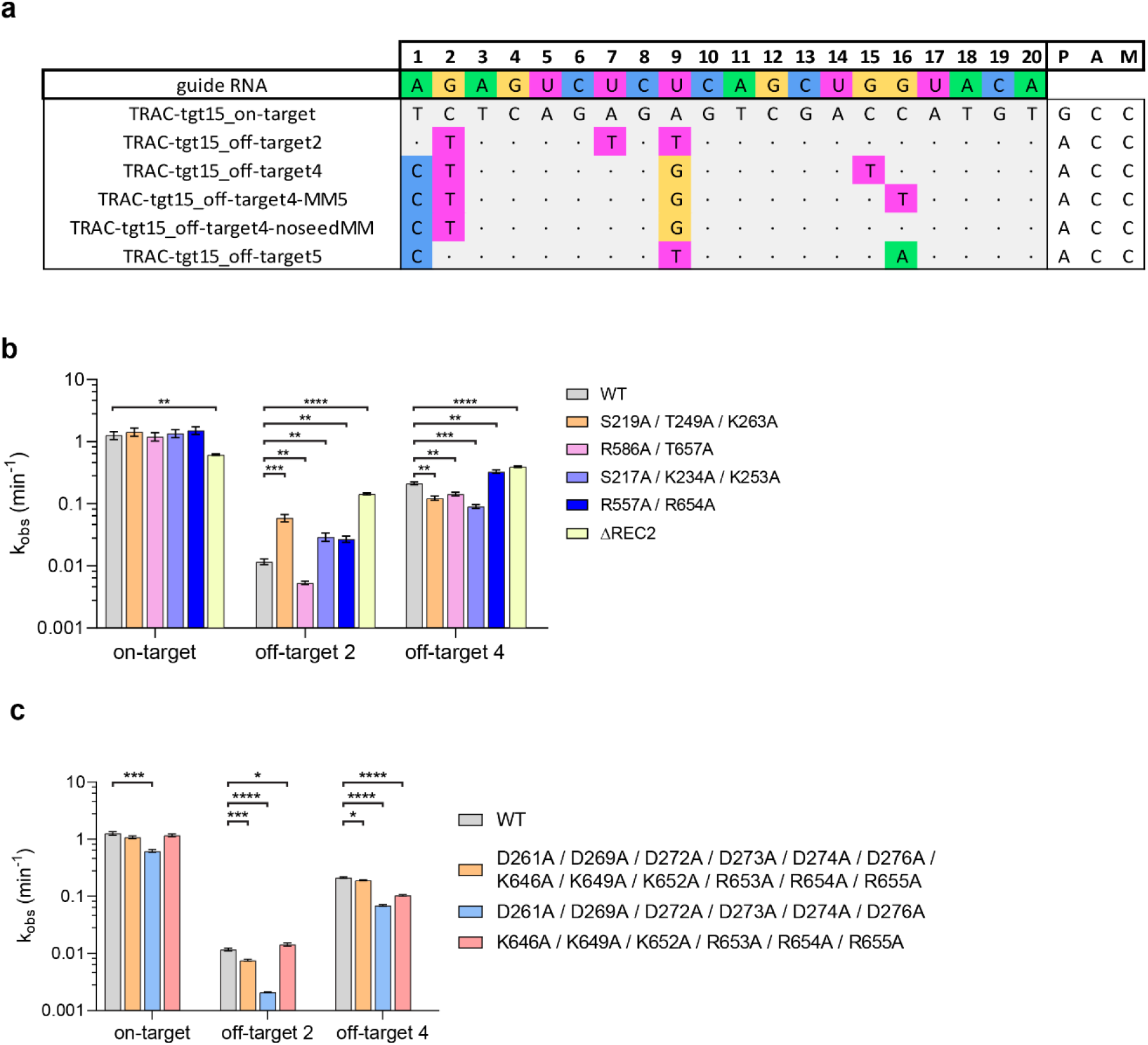
Examination of in vitro cleavage activity of structure guided REC2 and REC3 mutants of Cas9. **a**, Off-target sequences selected for nuclease activity assays. Nucleotide mismatches between the TRAC guide RNA and the target are highlighted; matching nucleotides are denoted by a dot. Data represents mean ± SEM (n=4), significance was determined by a two-tailed t-test, * p <0.05, ** = p <0.01, *** p <0.001, **** = p <0.0001. **b**, Cleavage rate constants of PAM-distal duplex stabilising REC2/REC3 mutants on on- and off-targets. **c**, Cleavage rate constants of DDD and RRR helix mutants of Cas9.

**Extended Data Figure 7.**
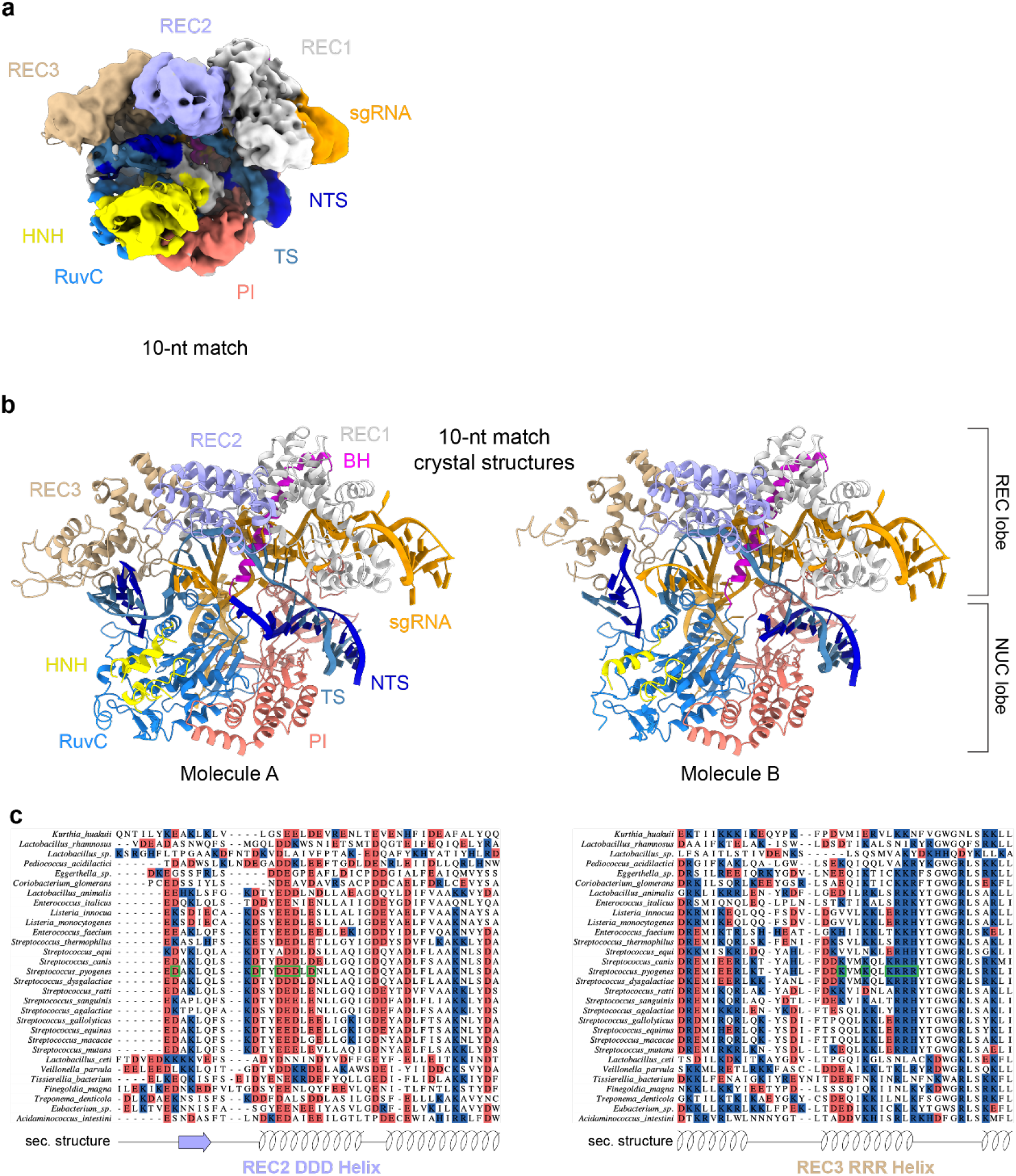
PAM-distal positioning and REC lobe conformation in the 10-nt match complex. **a**, CryoEM density of the 10-nt match complex overlaid with the structural model. NTS density can be traced along the heteroduplex (white). **b**, Cartoon representations of the X-ray crystallographic structures of the 10-nt match complex as based on the two complex copies (molecules A and B) in the crystallographic asymmetric unit. The complexes exhibit highly similar conformations (RMSD 0.46 Å). **c**, Alignment of protein sequences of the REC2 DDD and REC3 RRR helices from REC2-containing Cas9 orthologs.

**Extended Data Figure 8.**
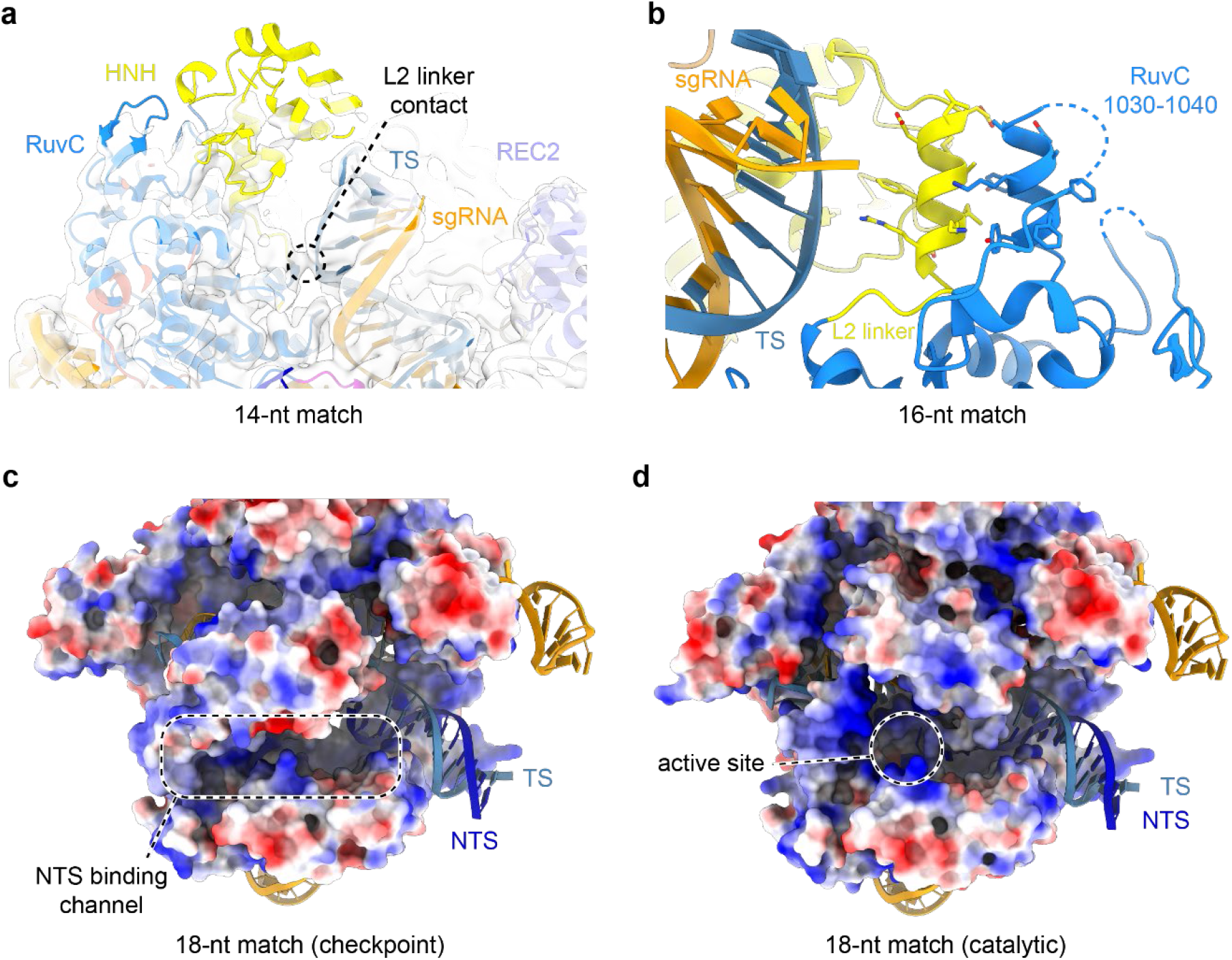
HNH undocking induced by R-loop extension. **a**, Residual HNH domain density (white) observed in the 14-nt match complex, in which the elongated heteroduplex establishes a contact with the L2 linker. No NTS density is observed past the PAM region due to disorder. **b**, Zoom-in view of the interaction between the HNH domain L2 linker and the RuvC 1030-1040 helix induced by heteroduplex proximity of the 16-bp complex. **c**, HNH domain relocation towards the binding channel results in the formation of a positively charged NTS binding channel. No residual electron density (white) is observed for the NTS in the absence of the PAM-distal duplex. The protein is coloured according to electrostatic surface potential, with red being negative, blue positive. **d**, Surface electrostatics map of the 18-nt match catalytic state of SpCas9, showing the NTS binding cleft with cleaved NTS positioned within the active site.

**Extended Data Figure 9.**
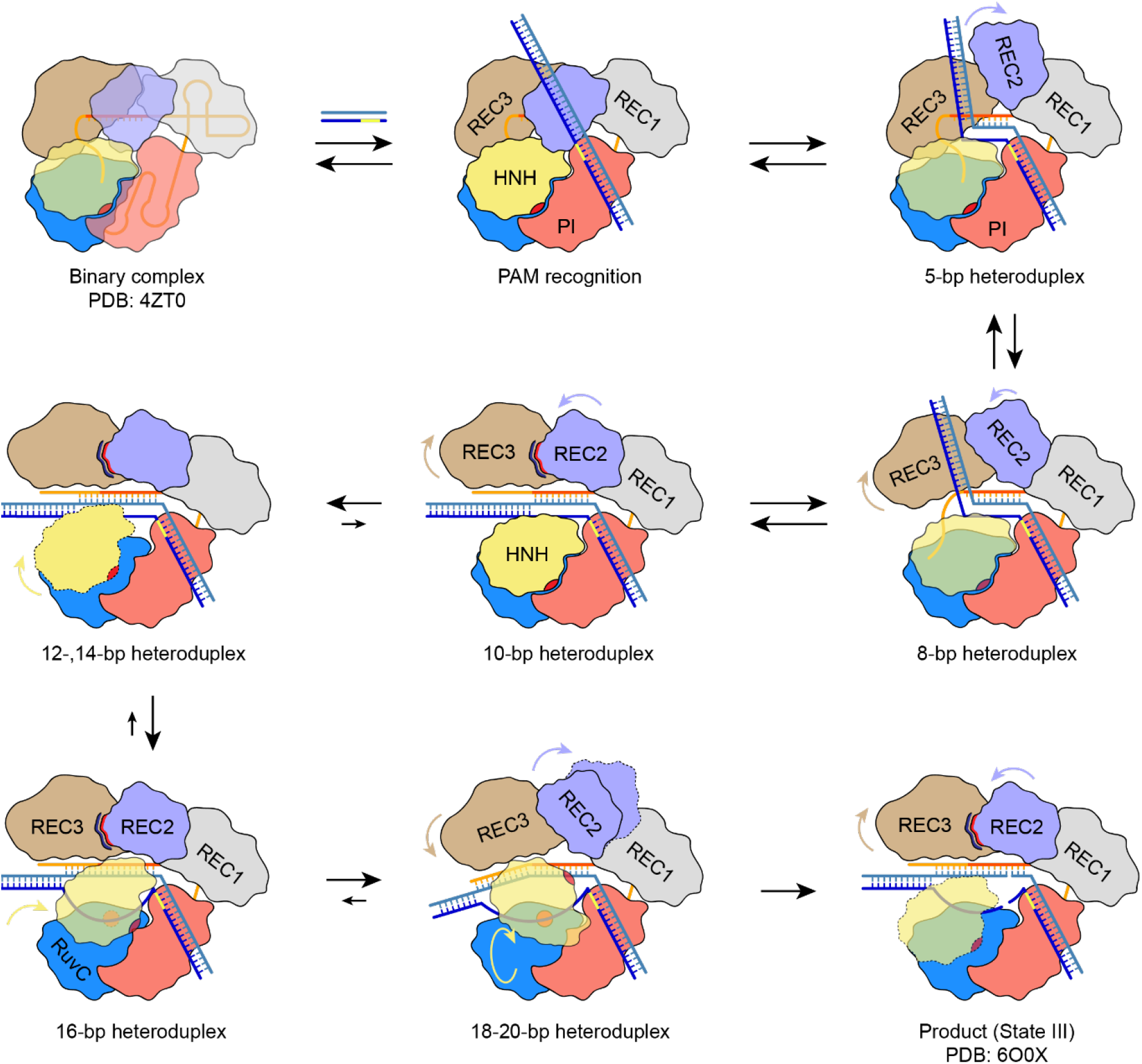
Molecular mechanism of Cas9 R-loop formation and conformational activation. In the RNA-bound (binary) complex, the central DNA binding channel is occluded by the REC2 and REC3 domains. Upon PAM recognition and initial 5-nt base pairing with the seed sequence of the guide RNA, the REC2 domain is displaced to form a binding cleft to accommodate the PAM-distal DNA duplex. Formation of 8-bp heteroduplex further displaces the REC3 domain and fully opens the central binding channel, while the PAM-distal duplex remains in the REC2/3 cavity. Extension of the R-loop to 10-bp heteroduplex places the guide-TS heteroduplex and the PAM-distal duplex into the central binding channel, accompanied by formation of electrostatic contacts between the REC2 and REC3 domains. Base pairing past the seed region results in undocking of the HNH domain from the RuvC and PI domain interface, and results in its repositioning towards target heteroduplex into the checkpoint state. R-loop formation past 18 base pairs induces REC2 domain displacement from the binding channel and rotation of the HNH domain active site towards the TS cleavage site, while simultaneously positioning the NTS in the RuvC domain active site.

**Extended Data Table 1.** Cryo-EM data collection, refinement and validation statistics

|  | dCas9-sgRNA (3-stem)-<br>6-nt match (PDB: 7Z4C,<br>EMDB: 14493) | dCas9-sgRNA (3-stem)-<br>8-nt match (PDB: 7Z4E,<br>EMDB: 14494) | dCas9-sgRNA (3-stem)-<br>10-nt match (PDB:<br>7Z4K, EMDB: 14500) |
| --- | --- | --- | --- |
| **Data collection and processing** |  |  |  |
| Magnification | 130,000 | 130,000 | 130,000 |
| Voltage (kV) | 300 | 300 | 300 |
| Electron exposure (e-/Å<sup>2</sup>) | 64.23 | 64.23 | 66.22 |
| Defocus range (μm) | -1.0 to -2.4 | -1.0 to -2.4 | -1.0 to -2.4 |
| Pixel size (Å) | 0.65 | 0.65 | 0.65 |
| Symmetry imposed | C1 | C1 | C1 |
| Initial particle images (no.) | 137,838 | 196,733 | 128,167 |
| Final particle images (no.) | 76,715 | 147,231 | 127,523 |
| Map resolution (Å) | 3.87 | 4.14 | 3.81 |
| FSC threshold | 0.143 | 0.143 | 0.143 |
| Map resolution range (Å) | 2.5-7.5 | 2.5-7.5 | 2.5-7.5 |
| **Refinement** |  |  |  |
| Initial model used (PDB code) | 5FQ5 | 5FQ5 | 5FQ5 |
| Model resolution (Å) | 4.2 | 4.5 | 4.3 |
| FSC threshold | 0.5 | 0.5 | 0.5 |
| Map sharpening *B* factor (Å<sup>2</sup>) | -164.1 | -208.6 | -141.8 |
| Model composition |  |  |  |
| Non-hydrogen atoms | 14180 | 14069 | 14005 |
| Protein residues | 1357 | 1357 | 1360 |
| Nucleotide residues | 148 | 142 | 138 |
| Ligands | 0 | 0 | 0 |
| *B* factors (mean, Å<sup>2</sup>) |  |  |  |
| Protein | 239.2 | 274.9 | 256.3 |
| Nucleotide | 190.1 | 212.3 | 196.6 |
| Ligand | n.a. | n.a. | n.a. |
| R.m.s. deviations |  |  |  |
| Bond lengths (Å) | 0.003 | 0.003 | 0.003 |
| Bond angles (°) | 0.451 | 0.546 | 0.436 |
| Validation |  |  |  |
| MolProbity score | 1.78 | 1.67 | 1.58 |
| Clashscore | 8.05 | 9.63 | 9.47 |
| Poor rotamers (%) | 2.06 | 1.64 | 0.99 |
| Ramachandran plot |  |  |  |
| Favored (%) | 97.5 | 98.0 | 97.6 |
| Allowed (%) | 2.5 | 2.0 | 2.4 |
| Disallowed (%) | 0.0 | 0.0 | 0.0 |

|  | dCas9-sgRNA (3-stem)-<br>12-nt match (PDB: 7Z4G,<br>EMDB: 14496) | dCas9-sgRNA (3-stem)-<br>14-nt match (PDB:<br>7Z4H, EMDB: 14497) | dCas9-sgRNA (3-stem)-<br>16-nt match (PDB: 7Z4I,<br>EMDB: 14498) |
| --- | --- | --- | --- |
| **Data collection and processing** |  |  |  |
| Magnification | 130,000 | 130,000 | 130,000 |
| Voltage (kV) | 300 | 300 | 300 |
| Electron exposure (e-/Å<sup>2</sup>) | 66.60 | 67.44 | 67.24 |
| Defocus range (μm) | -1.0 to -2.4 | -1.0 to -2.4 | -1.0 to -2.4 |
| Pixel size (Å) | 0.65 | 0.65 | 0.65 |
| Symmetry imposed | C1 | C1 | C1 |
| Initial particle images (no.) | 105,145 | 229,760 | 434,090 |
| Final particle images (no.) | 104,580 | 75,738 | 146,573 |
| Map resolution (Å) | 3.64 | 3.49 | 3.12 |
| FSC threshold | 0.143 | 0.143 | 0.143 |
| Map resolution range (Å) | 2.5-7.5 | 2.5-7.5 | 2.5-7.5 |
| **Refinement** |  |  |  |
| Initial model used (PDB code) | 5FQ5 | 5FQ5 | 5FQ5 |
| Model resolution (Å) | 3.9 | 3.8 | 3.4 |
| FSC threshold | 0.5 | 0.5 | 0.5 |
| Map sharpening *B* factor (Å<sup>2</sup>) | -122.8 | -117.1 | -98.9 |
| Model composition |  |  |  |
| Non-hydrogen atoms | 12714 | 13901 | 13834 |
| Protein residues | 1188 | 1334 | 1338 |
| Nucleotide residues | 144 | 143 | 137 |
| Ligands | 0 | 0 | 2 |
| *B* factors (mean, Å<sup>2</sup>) |  |  |  |
| Protein | 204.9 | 256.7 | 201.2 |
| Nucleotide | 187.3 | 196.9 | 138.1 |
| Ligand | n.a. | n.a. | 149.5 |
| R.m.s. deviations |  |  |  |
| Bond lengths (Å) | 0.003 | 0.003 | 0.003 |
| Bond angles (°) | 0.478 | 0.452 | 0.464 |
| Validation |  |  |  |
| MolProbity score | 1.63 | 1.60 | 1.43 |
| Clashscore | 12.71 | 9.94 | 7.91 |
| Poor rotamers (%) | 1.03 | 0.92 | 1.44 |
| Ramachandran plot |  |  |  |
| Favored (%) | 98.1 | 98.0 | 98.5 |
| Allowed (%) | 1.9 | 2.0 | 1.5 |
| Disallowed (%) | 0.0 | 0.0 | 0.0 |

|  | Cas9-sgRNA (3-stem)-<br>18-nt match (checkpoint)<br>(PDB: 7Z4L, EMDB:<br>14501) | Cas9-sgRNA (3-stem)-<br>18-nt match (catalytic)<br>(PDB: 7Z4J, EMDB:<br>14499) |
| --- | --- | --- |
| **Data collection and processing** |  |  |
| Magnification | 270,000 | 130,000 |
| Voltage (kV) | 300 | 300 |
| Electron exposure (e-/Å<sup>2</sup>) | 60.00 | 64.20 |
| Defocus range (μm) | -0.8 to -1.5 | -1.0 to -2.4 |
| Pixel size (Å) | 0.45 | 0.65 |
| Symmetry imposed | C1 | C1 |
| Initial particle images (no.) | 597,564 | 277,414 |
| Final particle images (no.) | 66,518 | 68,772 |
| Map resolution (Å) | 2.54 | 2.99 |
| FSC threshold | 0.143 | 0.143 |
| Map resolution range (Å) | 2.5-7.5 | 2.5-7.5 |
| **Refinement** |  |  |
| Initial model used (PDB code) | 5FQ5 | 5FQ5 |
| Model resolution (Å) | 2.9 | 3.2 |
| FSC threshold | 0.5 | 0.5 |
| Map sharpening *B* factor (Å<sup>2</sup>) | -48.1 | -75.8 |
| Model composition |  |  |
| Non-hydrogen atoms | 13822 | 13948 |
| Protein residues | 1337 | 1345 |
| Nucleotide residues | 136 | 142 |
| Ligands | 3 | 4 |
| *B* factors (mean, Å<sup>2</sup>) |  |  |
| Protein | 120.8 | 140.6 |
| Nucleotide | 78.7 | 121.4 |
| Ligand | 85.6 | 81.8 |
| R.m.s. deviations |  |  |
| Bond lengths (Å) | 0.005 | 0.005 |
| Bond angles (°) | 0.525 | 0.476 |
| Validation |  |  |
| MolProbity score | 1.88 | 1.69 |
| Clashscore | 8.74 | 7.21 |
| Poor rotamers (%) | 2.83 | 2.15 |
| Ramachandran plot |  |  |
| Favored (%) | 97.7 | 97.8 |
| Allowed (%) | 2.3 | 2.2 |
| Disallowed (%) | 0.0 | 0.0 |

**Extended Data Table 2.** Data collection and refinement statistics (molecular replacement)

|  | dCas9-sgRNA (2-stem)-<br>10-nt match (PDB: 7Z4D) |
| --- | --- |
| **Data collection** |  |
| Space group | P1 |
| Cell dimensions |  |
| *a*, *b*, *c* (Å) | 88.64, 95.51, 169.80 |
| $\alpha$, $\beta$, $\gamma$ (°) | 80.83, 78.04, 62.34 |
| Resolution (Å) | 66.49-3.10 (3.21 – 3.10) |
| *R*<sub>merge</sub> | 0.34 (2.18) |
| *I* / $\sigma I$ | 7.6 (1.3) |
| CC<sub>1/2</sub> | 0.988 (0.446) |
| Completeness (%) | 89.2 (56.7) |
| Multiplicity | 10.2 (10.1) |
| **Refinement** |  |
| No. reflections | 85696 (4963) |
| *R*<sub>work</sub> / *R*<sub>free</sub> | 24.7/27.5 |
| No. atoms |  |
| Macromolecules | 24978 |
| Ligands | 16 |
| Solvent | 88 |
| *B*-factors |  |
| Macromolecules | 69.4 |
| Ligands | 65.9 |
| Solvent | 29.7 |
| R.m.s. deviations |  |
| Bond lengths (Å) | 0.003 |
| Bond angles (°) | 0.560 |

\* Values in parentheses are for highest-resolution shell. Structure was solved from data collected with two identical crystals.

**Extended Data Table 3.**
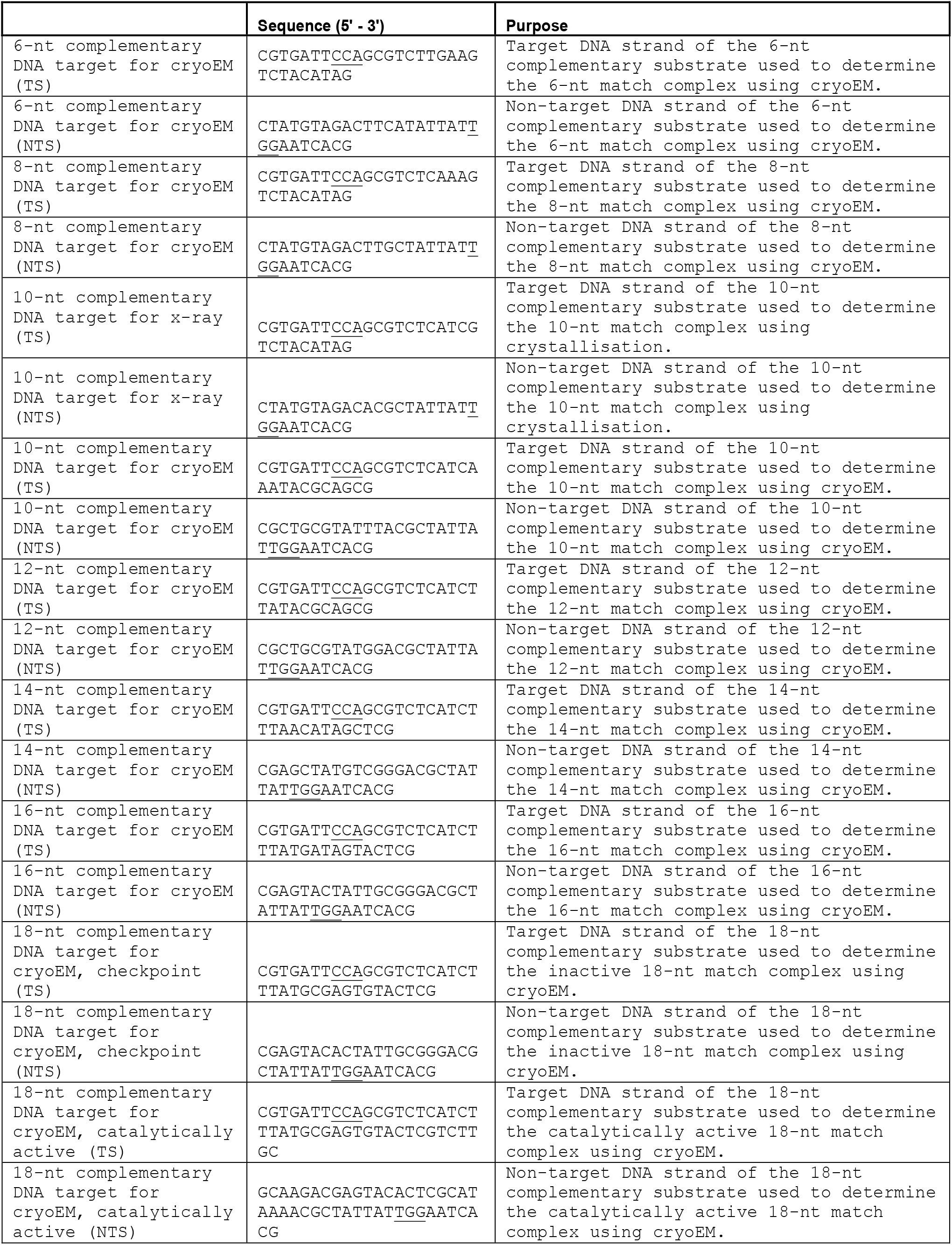
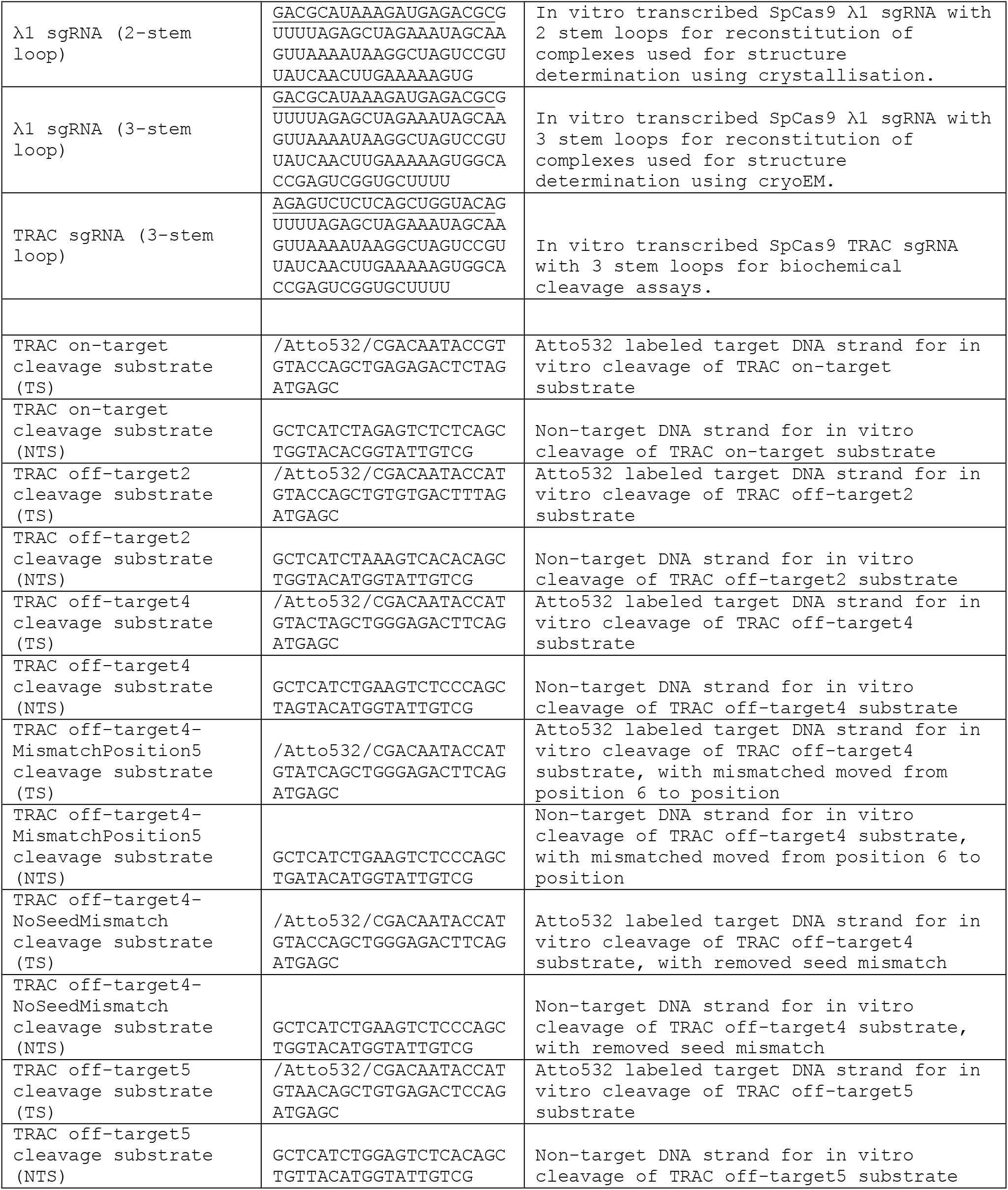
Oligonucleotide sequences used in the study. PAM sequence underlined.

## References

1 Garneau, J. E. et al. The CRISPR/Cas bacterial immune system cleaves bacteriophage and plasmid DNA. Nature 468, 67–71, doi:10.1038/nature09523 (2010).

2 Jinek, M. et al. A programmable dual-RNA-guided DNA endonuclease in adaptive bacterial immunity. Science 337, 816–821, doi:10.1126/science.1225829 (2012).

3 Sapranauskas, R. et al. The Streptococcus thermophilus CRISPR/Cas system provides immunity in Escherichia coli. Nucleic Acids Res 39, 9275–9282, doi:10.1093/nar/gkr606 (2011).

4 Cong, L. et al. Multiplex genome engineering using CRISPR/Cas systems. Science 339, 819–823, doi:10.1126/science.1231143 (2013).

5 Jinek, M. et al. RNA-programmed genome editing in human cells. Elife 2, e00471, doi:10.7554/eLife.00471 (2013).

6 Mali, P. et al. RNA-guided human genome engineering via Cas9. Science 339, 823–826, doi:10.1126/science.1232033 (2013).

7 Mekler, V., Minakhin, L. & Severinov, K. Mechanism of duplex DNA destabilization by RNA-guided Cas9 nuclease during target interrogation. Proc Natl Acad Sci U S A 114, 5443–5448, doi:10.1073/pnas.1619926114 (2017).

8 Sternberg, S. H., Redding, S., Jinek, M., Greene, E. C. & Doudna, J. A. DNA interrogation by the CRISPR RNA-guided endonuclease Cas9. Nature 507, 62–67, doi:10.1038/nature13011 (2014).

9 Szczelkun, M. D. et al. Direct observation of R-loop formation by single RNA-guided Cas9 and Cascade effector complexes. Proc Natl Acad Sci U S A 111, 9798–9803, doi:10.1073/pnas.1402597111 (2014).

10 Anders, C., Niewoehner, O., Duerst, A. & Jinek, M. Structural basis of PAM-dependent target DNA recognition by the Cas9 endonuclease. Nature 513, 569–573, doi:10.1038/nature13579 (2014).

11 Ivanov, I. E. et al. Cas9 interrogates DNA in discrete steps modulated by mismatches and supercoiling. Proc Natl Acad Sci U S A 117, 5853–5860, doi:10.1073/pnas.1913445117 (2020).

12 Jiang, F., Zhou, K., Ma, L., Gressel, S. & Doudna, J. A. STRUCTURAL BIOLOGY. A Cas9-guide RNA complex preorganized for target DNA recognition. Science 348, 1477–1481, doi:10.1126/science.aab1452 (2015).

13 Sternberg, S. H., LaFrance, B., Kaplan, M. & Doudna, J. A. Conformational control of DNA target cleavage by CRISPR-Cas9. Nature 527, 110–113, doi:10.1038/nature15544 (2015).

14 Cameron, P. et al. Mapping the genomic landscape of CRISPR-Cas9 cleavage. Nat Methods 14, 600–606, doi:10.1038/nmeth.4284 (2017).

15 Doench, J. G. et al. Optimized sgRNA design to maximize activity and minimize off-target effects of CRISPR-Cas9. Nat Biotechnol 34, 184–191, doi:10.1038/nbt.3437 (2016).

16 Hsu, P. D. et al. DNA targeting specificity of RNA-guided Cas9 nucleases. Nat Biotechnol 31, 827–832, doi:10.1038/nbt.2647 (2013).

17 Lazzarotto, C. R. et al. CHANGE-seq reveals genetic and epigenetic effects on CRISPR-Cas9 genome-wide activity. Nat Biotechnol, doi:10.1038/s41587-020-0555-7 (2020).

18 Tsai, S. Q. et al. GUIDE-seq enables genome-wide profiling of off-target cleavage by CRISPR-Cas nucleases. Nat Biotechnol 33, 187–197, doi:10.1038/nbt.3117 (2015).

19 Boyle, E. A. et al. Quantification of Cas9 binding and cleavage across diverse guide sequences maps landscapes of target engagement. Sci Adv 7, doi:10.1126/sciadv.abe5496 (2021).

20 Jones, S. K., Jr. et al. Massively parallel kinetic profiling of natural and engineered CRISPR nucleases. Nat Biotechnol, doi:10.1038/s41587-020-0646-5 (2020).

21 Zhang, L. et al. Systematic in vitro profiling of off-target affinity, cleavage and efficiency for CRISPR enzymes. Nucleic Acids Research, doi:10.1093/nar/gkaa231 (2020).

22 Singh, D., Sternberg, S. H., Fei, J., Doudna, J. A. & Ha, T. Real-time observation of DNA recognition and rejection by the RNA-guided endonuclease Cas9. Nat Commun 7, 12778, doi:10.1038/ncomms12778 (2016).

23 Chen, J. S. et al. Enhanced proofreading governs CRISPR-Cas9 targeting accuracy. Nature 550, 407–410, doi:10.1038/nature24268 (2017).

24 Dagdas, Y. S., Chen, J. S., Sternberg, S. H., Doudna, J. A. & Yildiz, A. A conformational checkpoint between DNA binding and cleavage by CRISPR-Cas9. Sci Adv 3, eaao0027, doi:10.1126/sciadv.aao0027 (2017).

25 Anders, C., Bargsten, K. & Jinek, M. Structural Plasticity of PAM Recognition by Engineered Variants of the RNA-Guided Endonuclease Cas9. Mol Cell 61, 895–902, doi:10.1016/j.molcel.2016.02.020 (2016).

26 Jiang, F. et al. Structures of a CRISPR-Cas9 R-loop complex primed for DNA cleavage. Science 351, 867–871, doi:10.1126/science.aad8282 (2016).

27 Nishimasu, H. et al. Crystal structure of Cas9 in complex with guide RNA and target DNA. Cell 156, 935–949, doi:10.1016/j.cell.2014.02.001 (2014).

28 Zhu, X. et al. Cryo-EM structures reveal coordinated domain motions that govern DNA cleavage by Cas9. Nat Struct Mol Biol 26, 679–685, doi:10.1038/s41594-019-0258-2 (2019).

29 Punjani, A. & Fleet, D. J. 3D variability analysis: Resolving continuous flexibility and discrete heterogeneity from single particle cryo-EM. J Struct Biol 213, 107702, doi:10.1016/j.jsb.2021.107702 (2021).

30 Cofsky, J. C., Soczek, K. M., Knott, G. J., Nogales, E. & Doudna, J. A. CRISPR-Cas9 bends and twists DNA to read its sequence. bioRxiv, 2021.2009.2006.459219, doi:10.1101/2021.09.06.459219 (2021).

31 Sung, K., Park, J., Kim, Y., Lee, N. K. & Kim, S. K. Target Specificity of Cas9 Nuclease via DNA Rearrangement Regulated by the REC2 Domain. J Am Chem Soc 140, 7778–7781, doi:10.1021/jacs.8b03102 (2018).

32 Yang, M. et al. The Conformational Dynamics of Cas9 Governing DNA Cleavage Are Revealed by Single-Molecule FRET. Cell Rep 22, 372–382, doi:10.1016/j.celrep.2017.12.048 (2018).

33 Sun, W. et al. Structures of Neisseria meningitidis Cas9 Complexes in Catalytically Poised and Anti-CRISPR-Inhibited States. Mol Cell 76, 938–952 e935, doi:10.1016/j.molcel.2019.09.025 (2019).

34 Zhang, Y. et al. Catalytic-state structure and engineering of Streptococcus thermophilus Cas9. Nature Catalysis 3, 813–823, doi:10.1038/s41929-020-00506-9 (2020).

35 Casalino, L., Nierzwicki, L., Jinek, M. & Palermo, G. Catalytic Mechanism of Non-Target DNA Cleavage in CRISPR-Cas9 Revealed by Ab Initio Molecular Dynamics. ACS Catal 10, 13596–13605, doi:10.1021/acscatal.0c03566 (2020).

36 Bravo, J. P. K. et al. Structural basis for mismatch surveillance by CRISPR/Cas9. bioRxiv, 2021.2009.2014.460224, doi:10.1101/2021.09.14.460224 (2021).

37 Klum, S. M., Chandradoss, S. D., Schirle, N. T., Joo, C. & MacRae, I. J. Helix-7 in Argonaute2 shapes the microRNA seed region for rapid target recognition. EMBO J 37, 75–88, doi:10.15252/embj.201796474 (2018).

38 Mulepati, S., Heroux, A. & Bailey, S. Structural biology. Crystal structure of a CRISPR RNA-guided surveillance complex bound to a ssDNA target. Science 345, 1479–1484, doi:10.1126/science.1256996 (2014).

39 Semenova, E. et al. Interference by clustered regularly interspaced short palindromic repeat (CRISPR) RNA is governed by a seed sequence. Proc Natl Acad Sci U S A 108, 10098–10103, doi:10.1073/pnas.1104144108 (2011).

40 Blosser, T. R. et al. Two distinct DNA binding modes guide dual roles of a CRISPR-Cas protein complex. Mol Cell 58, 60–70, doi:10.1016/j.molcel.2015.01.028 (2015).

41 Xiao, Y. et al. Structure Basis for Directional R-loop Formation and Substrate Handover Mechanisms in Type I CRISPR-Cas System. Cell 170, 48–60 e11, doi:10.1016/j.cell.2017.06.012 (2017).

42 Kuscu, C., Arslan, S., Singh, R., Thorpe, J. & Adli, M. Genome-wide analysis reveals characteristics of off-target sites bound by the Cas9 endonuclease. Nat Biotechnol 32, 677–683, doi:10.1038/nbt.2916 (2014).

43 Gaudelli, N. M. et al. Programmable base editing of A*T to G*C in genomic DNA without DNA cleavage. Nature 551, 464–471, doi:10.1038/nature24644 (2017).

44 Komor, A. C., Kim, Y. B., Packer, M. S., Zuris, J. A. & Liu, D. R. Programmable editing of a target base in genomic DNA without double-stranded DNA cleavage. Nature 533, 420–424, doi:10.1038/nature17946 (2016).

45 Maeder, M. L. et al. CRISPR RNA-guided activation of endogenous human genes. Nat Methods 10, 977–979, doi:10.1038/nmeth.2598 (2013).

46 Qi, L. S. et al. Repurposing CRISPR as an RNA-guided platform for sequence-specific control of gene expression. Cell 152, 1173–1183, doi:10.1016/j.cell.2013.02.022 (2013).

47 Anzalone, A. V., Koblan, L. W. & Liu, D. R. Genome editing with CRISPR-Cas nucleases, base editors, transposases and prime editors. Nat Biotechnol 38, 824–844, doi:10.1038/s41587-020-0561-9 (2020).

48 Pacesa, M. et al. Structural basis for Cas9 off-target activity. bioRxiv, 2021.2011.2018.469088, doi:10.1101/2021.11.18.469088 (2021).

49 Donohoue, P. D. et al. Conformational control of Cas9 by CRISPR hybrid RNA-DNA guides mitigates off-target activity in T cells. Mol Cell 81, 3637–3649 e3635, doi:10.1016/j.molcel.2021.07.035 (2021).

50 Newton, M. D. et al. DNA stretching induces Cas9 off-target activity. Nat Struct Mol Biol 26, 185–192, doi:10.1038/s41594-019-0188-z (2019).

51 Vonrhein, C. et al. Data processing and analysis with the autoPROC toolbox. Acta Crystallogr D Biol Crystallogr 67, 293–302, doi:10.1107/S0907444911007773 (2011).

52 Adams, P. D. et al. PHENIX: a comprehensive Python-based system for macromolecular structure solution. Acta Crystallogr D Biol Crystallogr 66, 213–221, doi:10.1107/S0907444909052925 (2010).

53 Punjani, A., Rubinstein, J. L., Fleet, D. J. & Brubaker, M. A. cryoSPARC: algorithms for rapid unsupervised cryo-EM structure determination. Nat Methods 14, 290–296, doi:10.1038/nmeth.4169 (2017).

54 Punjani, A., Zhang, H. & Fleet, D. J. Non-uniform refinement: adaptive regularization improves single-particle cryo-EM reconstruction. Nat Methods 17, 1214–1221, doi:10.1038/s41592-020-00990-8 (2020).

55 Pettersen, E. F. et al. UCSF ChimeraX: Structure visualization for researchers, educators, and developers. Protein Sci 30, 70–82, doi:10.1002/pro.3943 (2021).

56 Emsley, P., Lohkamp, B., Scott, W. G. & Cowtan, K. Features and development of Coot. Acta Crystallogr D Biol Crystallogr 66, 486–501, doi:10.1107/S0907444910007493 (2010).

57 Liebschner, D. et al. Macromolecular structure determination using X-rays, neutrons and electrons: recent developments in Phenix. Acta Crystallogr D Struct Biol 75, 861–877, doi:10.1107/S2059798319011471 (2019).

58 Afonine, P. V. et al. Real-space refinement in PHENIX for cryo-EM and crystallography. Acta Crystallogr D Struct Biol 74, 531–544, doi:10.1107/S2059798318006551 (2018).

59 Williams, C. J. et al. MolProbity: More and better reference data for improved all-atom structure validation. Protein Sci 27, 293–315, doi:10.1002/pro.3330 (2018).

60 Krissinel, E. & Henrick, K. Inference of macromolecular assemblies from crystalline state. J Mol Biol 372, 774–797, doi:10.1016/j.jmb.2007.05.022 (2007).

61 Li, S., Olson, W. K. & Lu, X. J. Web 3DNA 2.0 for the analysis, visualization, and modeling of 3D nucleic acid structures. Nucleic Acids Res 47, W26–W34, doi:10.1093/nar/gkz394 (2019).

62 UniProt, C. UniProt: the universal protein knowledgebase in 2021. Nucleic Acids Res 49, D480–D489, doi:10.1093/nar/gkaa1100 (2021).

63 Edgar, R. C. MUSCLE: multiple sequence alignment with high accuracy and high throughput. Nucleic Acids Res 32, 1792–1797, doi:10.1093/nar/gkh340 (2004).

64 Waterhouse, A. M., Procter, J. B., Martin, D. M., Clamp, M. & Barton, G. J. Jalview Version 2--a multiple sequence alignment editor and analysis workbench. Bioinformatics 25, 1189–1191, doi:10.1093/bioinformatics/btp033 (2009).

